# A fibroblast-centric network drives cold fibrosis in the tumor microenvironment of lung squamous cell carcinoma

**DOI:** 10.1101/2025.08.25.668405

**Authors:** Shoval Miyara, Shachaf Frenkel, Avi Mayo, Philippe Gascard, Michael Strasser, David Gibbs, Eviatar Weizman, Itay Ben Shalom, Yaniv Stein, Deng Pan, Joseph A. Caruso, Veena Sangwan, Nicholas Bertos, Julie Berube, Sophie Camilleri-Broet, Spyridon Oikonomopoulos, Haig Djambazian, Kfir Baruch Umansky, Jacob Elkahal, Shimrit Mayer, Pierre-Olivier Fiset, Jiannis Ragoussis, Miri Adler, Eldad Tzahor, Sui Huang, Lorenzo Ferri, Thea D. Tlsty, Ruth Scherz-Shouval, Uri Alon

## Abstract

The tumor microenvironment (TME) of chronic inflammation-associated cancers (CIACs) is shaped by cycles of injury and maladaptive repair, yet the principles organizing fibrotic stroma in these tumors remain unclear. Here, we applied the concept of hot versus cold fibrosis, originally credentialed in non-cancerous fibrosis of heart and kidney, to lung squamous cell carcinoma (LUSC), a prototypical CIAC. Single-cell transcriptomics of matched tumor and adjacent-normal tissue from 16 treatment-naive LUSC patients identified a cold fibrotic architecture in the LUSC TME: cancer-associated fibroblasts (CAFs) expanded and adopted myofibroblast and stress-response states, while macrophages were depleted. This macrophage-poor, CAF-rich stroma was maintained by CAF autocrine growth factor loops, including *TIMP1*, *INHBA*, *TGFB1*, and *GMFB*. In parallel, the immune compartment exhibited a hot tumor phenotype with abundant T and B cells, forming spatially distinct but molecularly engaged networks with CAFs. CAF gene programs typifying cold fibrosis in LUSC were conserved in other CIACs, including esophageal and gastric adenocarcinomas. These results redefine desmoplastic regions of tumors through the lens of a non-cancer fibrosis model, demonstrating that conserved stromal circuits constitute therapeutic vulnerabilities in CIACs.

## Introduction

Chronic inflammation-associated cancers (CIACs) are a class of tumors that arise in tissues undergoing chronic injury and maladaptive repair^1^. A prototypical example of CIAC is lung squamous cell carcinoma (LUSC). LUSC has limited therapeutic options and poor prognosis^2^ compared to lung adenocarcinoma (LUAD). Unlike LUAD, LUSC lacks clear oncogenic drivers and has proven largely refractory to targeted therapies^3,4^. Immunotherapies such as immune checkpoint blockade have offered some benefits, but durable long-term remissions remain uncommon^5^.

LUSC follows repeated airway epithelial damage, often triggered by smoking or pollution, which drives aberrant wound-healing processes that include inflammation, immune remodeling, and fibrosis^6^. These processes often give rise to ECM-rich tumor regions traditionally described as fibro-inflammatory or desmoplastic^7,8^, but can also be characterized as a fibrotic tumor microenvironment (TME) that actively supports tumor progression^9,10^. Molecular and cellular programs driving tissue repair and fibrotic TME in LUSC have been recently described^11^. However, there is a need for a better understanding of LUSC tumor-associated fibrosis, and potentially of CIAC-associated fibrosis, to devise strategies for treatment.

Here we apply an advance in modeling fibrosis in non-cancer contexts to the case of LUSC. We built on work on heart and kidney fibrosis based on a mathematical model of inflammation and fibrosis dynamics that predicted the emergence of two distinct immune-stromal states, referred to as hot and cold fibrosis ^12–15^. This model, grounded in *in vitro* co-culture experiments^16^, suggests that hot fibrosis arises when macrophages and activated fibroblasts (named myofibroblasts) mutually support each other’s growth by reciprocal growth factor exchange. In contrast cold fibrosis is characterized by self-sustaining myofibroblasts with minimal macrophage involvement. Consistent with these predictions, we and others have shown that fibrotic tissues can adopt distinct hot and cold fibrosis configurations *in vivo*, with implications for treatment^14,15^. In particular, targeting the autocrine loop that sustains myofibroblasts can prevent and reverse fibrosis in the injured mouse heart and liver^14,17^. It remains unclear how these non-cancerous fibrotic states relate to the fibrotic-like TME of CIACs such as LUSC.

To avoid confusion, we note that the terminology of cold and hot fibrosis is specific to the local neighborhood of fibroblasts - hot fibrosis has macrophages adjacent to fibroblasts, whereas cold fibrosis does not. This terminology was inspired by the widely used concept of a hot and cold tumor, which refers to presence or absence of immune cells, including lymphocytes in the TME. Thus, one can observe a *hot tumor associated with cold fibrosis*, for example desmoplastic regions with numerous cancer-associated fibroblasts (CAFs) but rare macrophages, adjacent to regions rich with T and B cells.

Here, we test these concepts using LUSC as a model CIAC. We generated a single-cell transcriptomic atlas from 16 treatment-naive LUSC patients, profiling matched tumor and adjacent histologically-normal (referred to as adjacent-normal in the following) lung tissues. Using Pareto task inference (ParTI)^18^ we uncover how epithelial cells and CAFs shift their functional programs in the LUSC TME - epithelial cells acquire squamous cell identity and immune interaction expression programs, whearas fibroblasts acquire cellular stress and myofibroblastic CAF (myCAF)-like phenotypes. We then applied the conceptual frameworks of hot vs. cold fibrosis and tumor immunogenicity to classify the TME along two axes: adaptive immune infiltration and fibrotic state (hot or cold). We find that LUSC tumors are characterized by a dual phenotype of cold fibrotic TME, marked by abundant CAFs with macrophage depletion, and a hot tumor defined by increased T and B-cell infiltration. By integrating transcriptomic data with hierarchical analysis of intercellular networks, we show that CAFs emerge as central signaling hubs in the LUSC TME, with a prominent autocrine growth factor loop. This network structure contrasts with non-CIAC tumor types, such as breast cancer, where reciprocal macrophage-CAF interactions dominate and represent a hot fibrotic TME architecture ^19^. This study provides a framework for interpreting stromal and immune organization in CIACs and identifies potential cellular targets for modulating the fibrotic and immune components of the LUSC niche.

## Results

### LUSC tumor microenvironment shows cold fibrosis and hot tumor characteristics

To analyze the LUSC TME we recruited a prospective cohort of treatment-naive LUSC patients (n=16) and obtained matched LUSC tumor and adjacent-normal lung tissue samples from each patient at surgery. The adjacent-normal samples showed a classical lung histology, whereas the tumor samples showed squamous epithelial regions adjacent to desmoplastic (fibrotic) regions (Figure 1A).

**Figure 1.**
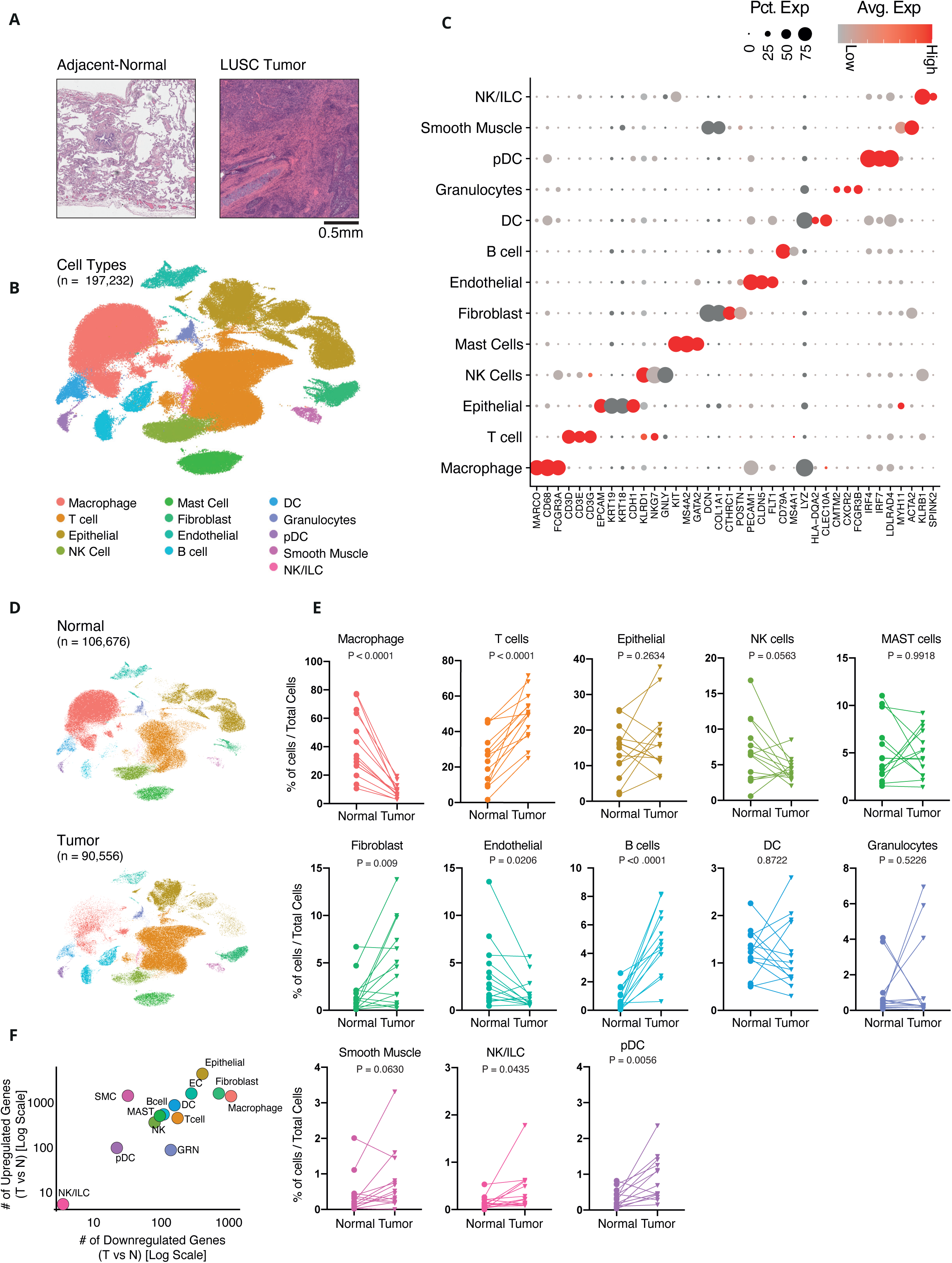
An atlas of the LUSC TME reveals a cold fibrosis-hot tumor phenotype. **A.** Representative hematoxylin and eosin (H&E) stained images of matched adjacent-normal lung tissue (left) and lung squamous cell carcinoma (LUSC) tumor tissue (right) from the same patient. Scale bar, 0.5mm. **B.** Uniform manifold approximation and projection (UMAP) embedding of 197,232 cells and delineation of 13 cell types including Macrophage, T cells, Epithelial Cells (including cancer epithelial cells), Natural Killer (NK) cells, Mast cells, Fibroblasts, Endothelial Cells, B Cells, Dendritic Cells (DC), Granulocytes, Plasmacytoid Dendritic Cells (pDC), Smooth Muscle cells, and NK/Innate Lymphoid Cells (ILC). Individual cells are colored by cell type. **C.** Dot plot showing the expression of marker genes across major cell types in the LUSC tumor and adjacent-normal lung microenvironment. Each row represents a cell type, and each column corresponds to a selected marker gene. The size of each dot indicates the percentage of cells within a given cell type expressing the gene (Pct. Exp), while the color scale represents the average expression level (Avg. Exp), from low (gray) to high (red). **D**.UMAP visualization of single-cell transcriptomes from adjacent-normal lung tissue (top; n = 106,676 cells) and LUSC tumor tissue (bottom; n = 90,556 cells). Individual cells are colored by cell type. **E.** Comparison of cell type abundances between adjacent-normal and LUSC tumor tissues on a per-patient basis (n = 14 patients with paired tumor and adjacent-normal samples). Each plot shows the percentage of a given cell type out of total cells in matched adjacent-normal and tumor samples, with paired lines representing individual patients. Statistical significance was calculated using a paired two-tailed Student’s t-test. **F.** Quantification of differentially expressed genes across major cell types between tumor (T) and adjacent-normal (N) lung tissue. Each point represents a cell type, plotted by the number of genes upregulated (y-axis) versus downregulated (x-axis) in the tumor relative to adjacent-normal tissue, both on a log_10_ scale.

We performed scRNA-sequencing of these samples using the 10x Genomics platform (Methods). After quality control and preprocessing, the dataset comprised 197,232 cells, including 90,556 tumor-derived and 106,676 adjacent-normal cells (Methods, Figure 1B). Unsupervised clustering and marker gene analysis identified 13 distinct cell types (Figure 1C, Methods, Table S1). These cell populations included macrophages (*MARCO, CD68*), T cells (*CD3D, CD3E*), epithelial cells, including cancer epithelial cells (*EPCAM, CDH1*), natural killer (NK) cells (*NKG7, GNLY*), mast cells (*KIT, MS4A2)*, fibroblasts (*DCN, COL1A1*), endothelial cells (*PECAM1, CLDN5*), B cells (*CD79A, MS4A1*), dendritic cells (DCs) (*HLA-DQA2, CLEC10A*), granulocytes (*CMTM2, CXCR2)*, plasmacytoid dendritic cells (pDCs) (*IRF7, IRF4*), smooth muscle cells (*MYH11, ACTA2*), and innate lymphoid cells (ILCs) (*SPINK2, KLRB1*) (Figure 1C, Table S1, Methods).

The distribution of these cell types differed between tumor and adjacent-normal tissues (Figure 1D). Several populations were present at comparable levels in tumor and adjacent-normal tissues including epithelial cells and innate immune subsets (NK cells, granulocytes, DCs, and mast cells). However, other populations exhibited consistent differences between the TME and adjacent-normal tissues.

Macrophages were depleted in tumor samples, showing an average 4.6-fold reduction across patients (*p* < 0.0001) (Figure 1E). In contrast, fibroblasts (CAFs) were enriched in tumor samples, with an average 2.9-fold increase (*p* < 0.009) (Figure 1E). CAF expansion and macrophage depletion are consistent with a cold fibrotic TME, in contrast to hot fibrosis, which is defined by abundant CAF and macrophage infiltration^12,14,19^.

Adaptive immune cells also demonstrated marked differences between tumor and adjacent-normal tissues. Both T and B-cells were increased in tumor samples relative to adjacent-normal tissue (*p* < 0.0001), consistent with a “hot” TME^20,21^. These observations indicate that the LUSC TME exhibits a dual phenotype, i.e. a cold fibrosis state alongside a hot tumor infiltration of adaptive immune components. As mentioned in the introduction, we note that cold fibrosis is defined by the absence of macrophages specifically, and thus a tumor can be hot (associated with many lymphocytes) but its fibrotic state can be cold (lack of macrophages). Importantly, spatial analysis of the LUSC TME revealed that T cells and CAFs tend to occupy distinct regions and therefore exhibit a spatial pattern of mutual avoidance^22^.

To further characterize the TME, we performed differential gene expression (DEG) analysis across tumor and adjacent-normal samples. The most transcriptionally active cell types included epithelial cells, as well as fibroblasts, macrophages, and endothelial cells (Figure 1F, Figure S1A, Table S2). As expected tumor epithelial cells showed increased cellular proliferation (Figure S1B).

These data support a model in which both fibrosis and immune infiltration shape the LUSC TME. We identified a cold fibrotic stroma marked by CAF expansion and macrophage exclusion, coexisting with a hot adaptive immune infiltration characterized by proliferative cancer epithelial cells and an enrichment in T and B cells.

### Epithelial cells in LUSC shift their division of labor into a squamous epithelial phenotype associated with increased translation, inflammation, and immune interaction

Next we analyzed the transcriptional program of single cell populations. Seurat analysis of LUSC and adjacent-normal epithelial cells revealed 13 distinct clusters, some of which were unique to tumor-derived cells (Figure 2A-C, Figure S2A, Table S3, Methods). To dissect the functional heterogeneity within the epithelium, we applied ParTI to the single-cell expression data^18^. ParTI analyzes a continuum of gene expression, capturing tradeoffs between specialized functions within a cell type. It identifies whether a population of cells can be approximated by a convex polytope (triangle, tetrahedron, and so on) in gene expression space. The vertices of this polytope, referred to as archetypes, represent transcriptional programs associated with task specialization. Cells near the center of the polytope co-express multiple programs and are considered generalists, while cells near the archetypes specialize in distinct biological functions. In epithelial cells, representing the cancer cell compartment in LUSC, ParTI identified a tetrahedral structure in gene expression space (*p* < 10⁻⁵), consistent with four archetypes (Figure 2D). GO analysis of genes enriched near each vertex revealed that the archetypes represent discrete biological programs (Figure 2E, Table S4). Archetype 1 was defined by high expression of *TP63*, a master regulator of squamous cell carcinomas^23,24^, and was enriched for genes involved in translation and NF-κB signaling. Archetype 2 was characterized by elevated expression of *MUC5B* and genes related to inflammation and IL-1 signaling, consistent with a pro-inflammatory epithelial phenotype. In contrast, Archetype 3 expressed *SFTPA2*, a hallmark of surfactant production and alveolar type II cell identity^25^, indicating a role in lung homeostasis. Archetype 4 was marked by high *TEKT1* expression, with GO term analysis reflecting a columnar ciliated epithelial phenotype involved in mucociliary clearance and epithelial barrier function^26^ (Figure 2E-F).

**Figure 2.**
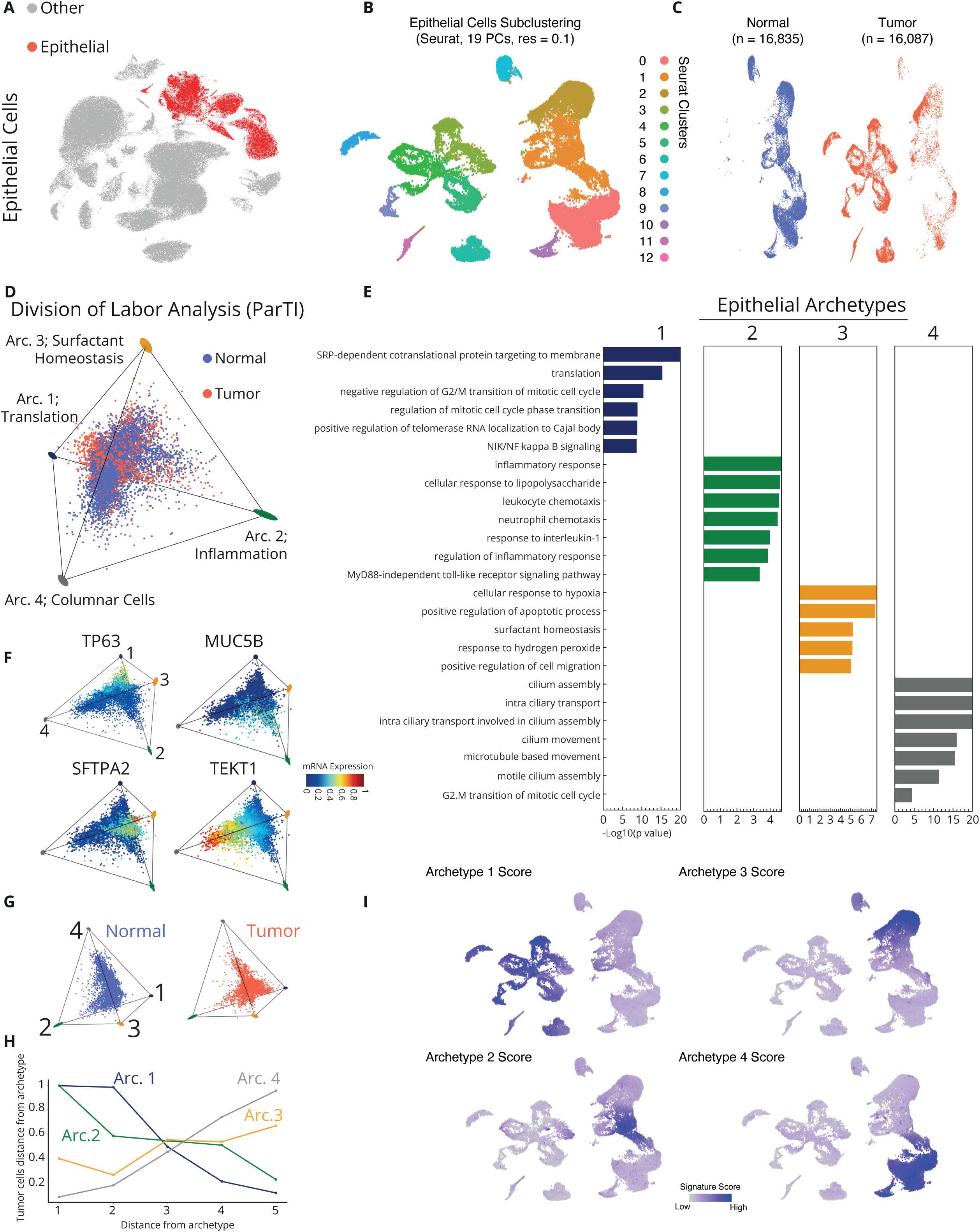
LUSC, associated epithelial cells acquire distinct archetypes characterized by enhanced translation, inflammation, and immune interaction. **A.** UMAP embedding of all cells, with epithelial cells highlighted in red and all other cell types shown in gray. **B.** UMAP plot of epithelial cells only, following subclustering using Seurat (19 principal components, resolution = 0.1), identifying 13 transcriptionally distinct epithelial subpopulations (clusters 0-12). **C.** Same UMAP as in (B), now split by condition, showing epithelial cells derived from adjacent-normal (left, n = 16,835) and tumor (right, n = 16,087) tissues. **D.** Pareto Task Inference (ParTI)^18^ was used to characterize the continuum of gene expression programs within epithelial cells across tumor and adjacent-normal tissues. Cells were projected into gene expression space and approximated by a tetrahedron *(p < 10*⁻⁵), indicating four transcriptional archetypes. Cells are projected on the first three principal components. **E.** Bar plot showing the top enriched Gene Ontology (GO) terms for each epithelial archetype (1-4) identified by ParTI analysis. GO terms are ranked by statistical significance (-log₁₀ p-value). **F.** Examples of top marker genes associated with each epithelial archetype (Table S4). The expression of *TP63* (archetype 1), *MUC5B* (archetype 2), *SFTPA2* (archetype 3), and *TEKT1* (archetype 4) is shown. Each gene’s expression is overlaid onto the tetrahedral projection of epithelial cells from ParTI analysis, where cells are colored by normalized mRNA expression (blue: low, red: high). **G.** Projection of adjacent-normal (left, blue) and tumor-derived (right, red) epithelial cells onto the tetrahedral archetype structure identified by ParTI. Each dot represents a single cell, positioned in gene expression space relative to the four identified archetypes. **H.** Enrichment of tumor-derived epithelial cells near each archetype, plotted as a function of Euclidean distance from the archetype vertex. **I.** Gene signature enrichment scores for each of the four epithelial archetypes (1-4) were calculated and projected onto a UMAP of epithelial cells. Cells are colored by signature score (purple to dark blue; linear scale).

Epithelial cells from adjacent-normal lung tissue were predominantly associated with the surfactant-producing and ciliated archetypes (archetypes 3 and 4), consistent with their homeostatic functions (Figure 2G-H). In contrast, archetype 1 (squamous epithelial program) and archetype 2 (inflammation program) were exclusively enriched in tumor-derived epithelium (Figure 2G-I). This redistribution of archetypes paralleled global changes in epithelial gene expression and proliferation (Figure 1E-F, Figure S1B), highlighting a shift from a homeostatic, columnar epithelial identity, supporting gas exchange and mucociliary clearance, toward a proliferative, *TP63*⁺ squamous program characteristic of LUSC.

### LUSC-associated fibroblasts adopt distinct archetypes associated with myofibroblast and stress-response phenotypes

Fibroblasts showed 6 clusters with a sharp division between tumor and adjacent-normal derived cells (Figure 3A-B, Figure S3A-B, Table S5, Methods). Four of these clusters (#0, 2, 3, 4) were similar to a myCAF state^27^ (*POSTN, ACAT2, MMP11*), cluster #3 showed similarties to an inflammatory CAF (iCAF) state (*CXCL6, CXCL12*)^27^, and cluster #1 to tissue-resident fibroblasts (*PI16*)^27,28^ (Figure S3B, D).

**Figure 3.**
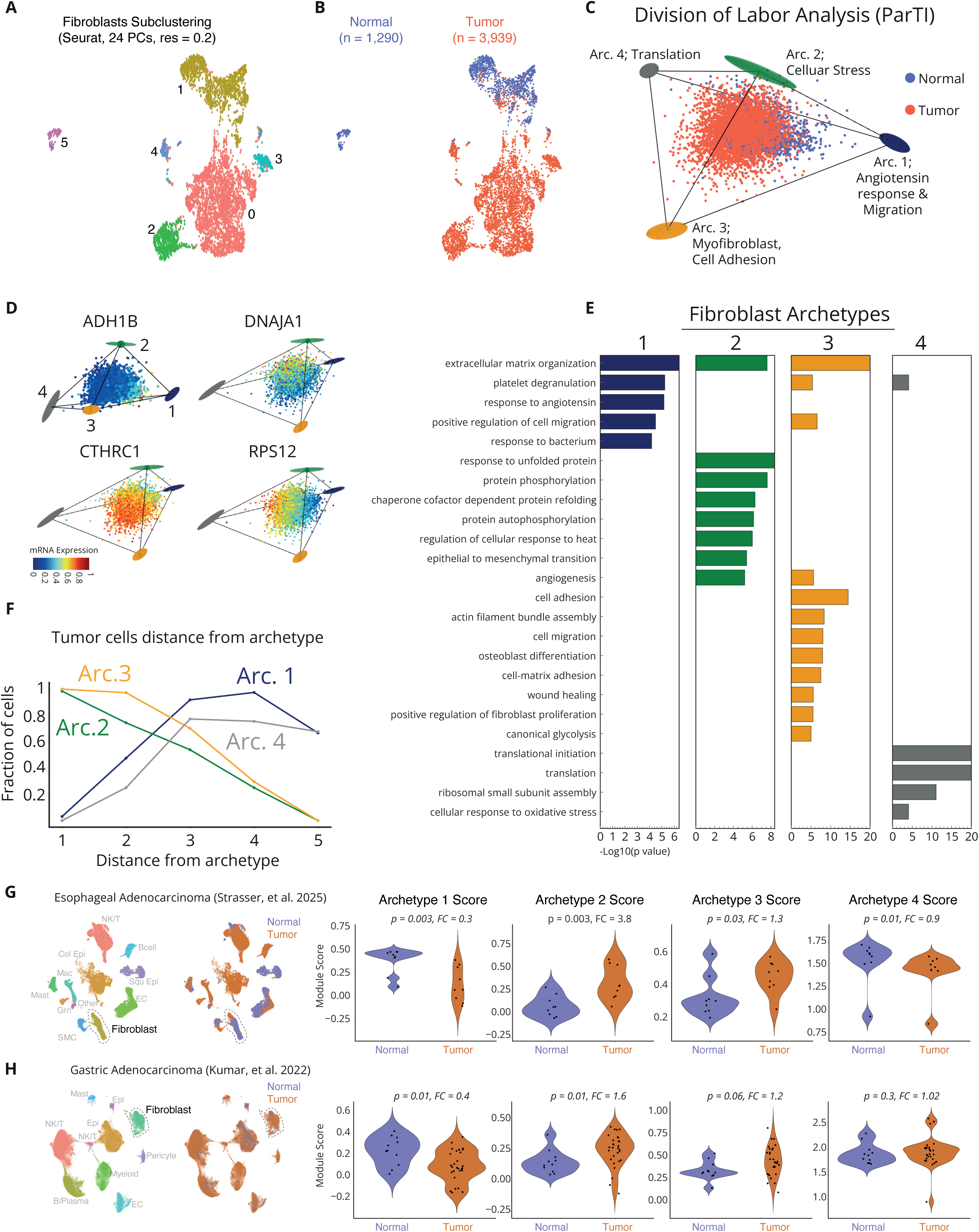
LUSC fibroblasts acquire two distinct archetypes are acquired, associated with a myofibroblast phenotype and stress response. **A.** UMAP embedding of all fibroblasts, with six transcriptionally distinct subpopulations identified by Seurat subclustering (24 principal components, resolution = 0.2; clusters 0–5). **B.** Same UMAP as in (A), now colored by tissue of origin, showing fibroblasts derived from adjacent-normal lung (blue, n = 1,290) and LUSC tumor tissue (red, n = 3,939). **C.** Pareto Task Inference (ParTI)^18^ was used to characterize the continuum of gene expression programs within fibroblasts from adjacent-normal and tumor samples. Cells were projected into gene expression space and approximated by a tetrahedron *(p = 0.03)*, indicative of four transcriptional archetypes. Cells are projected onto the first three principal components. **D.** Examples of top marker genes associated with each fibroblast archetype (Table S6). The expression of *ADH1B* (archetype 1), *DNAJA1* (archetype 2), CTHRC1 (archetype 3), and *RPS12* (archetype 4) is shown. Each gene’s expression is overlaid onto the tetrahedral projection of fibroblasts from ParTI analysis, where cells are colored by normalized mRNA expression (blue: low, red: high). **E.** Bar plot showing the top enriched Gene Ontology (GO) terms for each fibroblast archetype (1-4) identified by ParTI analysis. GO terms are ranked by statistical significance (-log₁₀ p-value). **F.** Enrichment of tumor-derived fibroblasts near each archetype, plotted as a function of Euclidean distance from the archetype vertex. **G.** left-UMAP embedding of single-cell RNA-seq data from esophageal adenocarcinoma (EAC) samples^33^, with cells colored by tissue origin (normal: blue; tumor: orange). Fibroblasts are outlined. Right-Violin plots show gene signature scores for each fibroblast archetype (1-4) in normal versus tumor-derived fibroblasts, per patient. Each dot represents a biological replicate. Each plot displays the paired Wilcoxon p-value and the corresponding fold-change (tumor vs. normal) for the respective fibroblast archetype. **H.** Left, UMAP embedding of single-cell RNA-seq data from gastric adenocarcinoma (GAC)^34^, with cells colored by tissue origin (normal: blue; tumor: orange). The fibroblast cluster is outlined. Right, Violin plots show gene signature scores for each fibroblast archetype (1-4) in normal versus tumor-derived fibroblasts, per patient. Each dot represents a biological replicate. Each plot displays the paired Wilcoxon p-value and the corresponding fold-change (tumor vs. normal) for the respective fibroblast archetype.

To characterize fibroblasts’ functional division of labor in the LUSC TME, we applied ParTI to fibroblast populations from tumor and adjacent-normal tissues (Figure 3C). This analysis revealed a tetrahedral structure *(p = 0.03)*, indicative of four major transcriptional programs (Figure 3D-E, Table S6). Archetype 1 was marked by high expression of *ADH1B* and enriched for genes involved in angiotensin signaling and cell migration, consistent with a migratory or vascular-interacting fibroblast phenotype. Archetype 2 expressed high levels of the stress related protein DNAJA1^29,30^, and was enriched for GO terms involved in angiogenesis and the unfolded protein response, suggestive of a stress-adapted CAF phenotype. Archetype 3 showed elevated expression of *CTHRC1*, *POSTN,* and *ACTA2*, conserved markers of activated myofibroblasts across tissues and disease contexts^27,28^, and was enriched for gene programs related to wound healing, ECM remodeling, and osteoblast differentiation (Figure 3D-E, Table S6). Archetype 4 was defined by high expression of ribosomal genes such as *RPS12*, indicating a fibroblast state with elevated translational activity.

Analysis of cells near the vertices revealed that fibroblasts from adjacent-normal lung tissues were primarily localized near the translation- and angiotensin-associated archetypes (archetypes 1 and 4), indicative of a homeostatic or quiescent fibroblast population (Figure 3F, Figure S3C). In contrast, CAFs were predominantly distributed near archetypes 2 and 3, reflecting transcriptional programs linked to stress adaptation, matrix production, and contractility (Figure 3F, Figure S3C). This shift suggests that CAFs in the LUSC TME undergo functional reprogramming toward an activated, myofibroblast state, with subsets acquiring features consistent with cellular stress. Scoring Seurat clusters by archetype-specific gene signatures corroborated the adjacent-normal (Archetype 1) and tumor enriched (Archetype 2 and 3) archetypes identified by ParTI analysis. Notably, archetype 4 was present across all Seurat clusters, representing a functional state not captured by clustering alone and is a case where ParTI provides information that is not visible in cluster analysis (Figure S3E).

To assess whether the fibroblast archetypes identified in LUSC are unique to this tumor or represent a broader response conserved across CIACs, we analyzed fibroblast archetype gene signatures in two additional CIACs: Esophageal adenocarcinoma (EAC), which arises from Barrett’s esophagus due to chronic acid reflux^31^, and gastric adenocarcinoma (GAC), a subset of which is driven by chronic inflammation associated with *Helicobacter pylori* infection^32^. Using two published single-cell RNA-seq datasets^33,34^, we calculated archetype gene signature scores in fibroblasts from normal and tumor tissues (Figure 3G-H). Archetype 1, associated with angiotensin signaling and quiescent fibroblasts, was markedly reduced in tumor tissues compared to normal fibroblasts in both EAC (70% reduction) and GAC (60% reduction). In contrast, tumor-enriched archetypes 2 and 3, linked to cellular stress and myofibroblast ECM remodeling, respectively, were significantly increased in both cancer types (Figure 3G-H). Specifically, Archetype 2 and 3 scores were elevated 3.8-fold in EAC and 1.6-fold in GAC CAFs, respectively.

We conclude that functional reprogramming of CAFs toward stress-adapted and myofibroblast states is a conserved feature of the TME across diverse CIACs, reflecting a shared division of labor and potential common therapeutic vulnerabilities.

### Network analysis reveals that the LUSC TME has features of concomitant cold fibrosis and T-cell engagement

the macrophage-fibroblast cell-circuit model underlying cold fibrosis is built on reciprocal growth factor exchange sustaining both cell types^12,14,16^. To explore cell-cell communication in the LUSC TME, we inferred interactions mediated by secreted growth-factor ligands using NicheNet, which integrates transcriptomic data with known ligand-receptor interactions and downstream signaling targets^35^. To specifically analyze growth factor-mediated interactions, we filtered ligands based on a curated list of known growth factors (Methods). We compared the growth factor signaling networks of tumor and adjacent-normal tissues based on each cell type outgoing and incoming signaling interactions, directionality, and cell abundance (Figure 4A, Methods).

**Figure 4.**
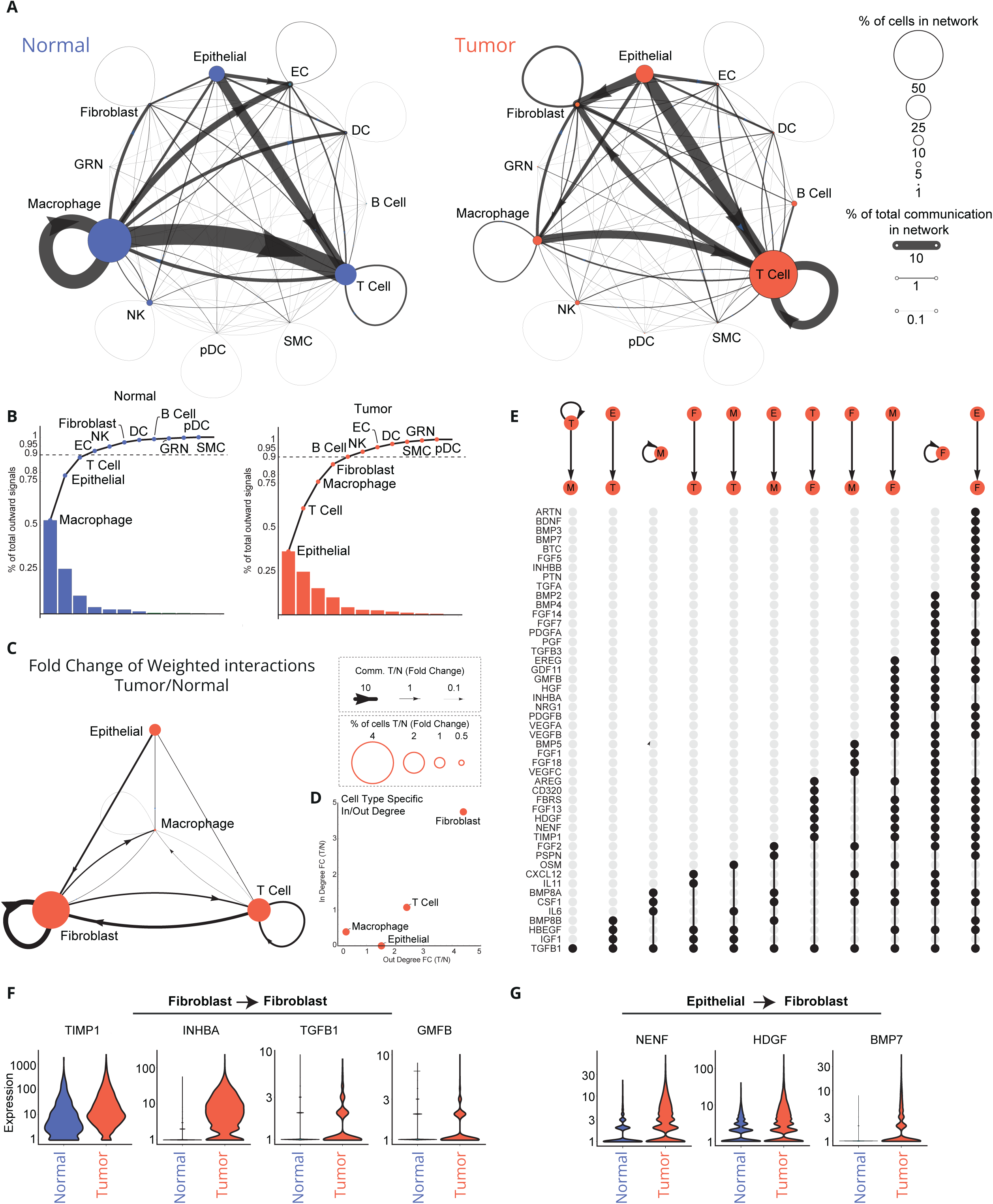
Network of growth factor interactions in tumor and adjacent-normal LUSC. **A.** Network of inferred growth factor–mediated interactions in adjacent-normal (left, blue) and tumor (right, red) tissue, as predicted by NicheNet. Nodes represent major cell types and are sized according to their relative abundance in the dataset (% of total cells). Directed edges represent ligand-receptor-mediated communication between sender and receiver cell types. Edge thickness reflects the relative contribution of each interaction to the total signaling within the network (% of total predicted communication). **B.** Fraction of outgoing growth factor interactions contributed by each cell type in adjacent-normal (left, blue) and LUSC tumor (right, red) samples. Bars represent the proportion of total outgoing signals attributed to each cell type, sorted in descending order. The overlaid line shows the cumulative contribution across cell types, with horizontal dashed lines marking the 90% cumulative thresholds. **C.** Fold-change network visualization comparing growth factor–mediated interactions in LUSC tumor versus adjacent-normal tissue. Nodes represent cell types and are scaled by the fold change in relative abundance (tumor/adjacent-normal). Directed edges represent ligand-receptor interactions, with edge thickness proportional to the fold change in interaction strength (tumor/adjacent-normal). Only interactions contributing to the top 90% of total network communication are shown. **D.** Scatter plot showing the fold change in incoming (y-axis) versus outgoing (x-axis) signaling for each cell type. **E.** Inferred ligand contributions to cell–cell interactions within the fold-change network of LUSC tumor versus adjacent-normal tissue. Arrows above each column represent directional communication between the sender (top node) and receiver (bottom node) cell types, including epithelial cells (E), fibroblasts (F), T cells (T), and macrophages (M). Each dot indicates a predicted ligand mediating the interaction, as inferred by NicheNet. **F.** Expression of *TIMP1*, *INHBA*, *TGFB1*, and *GMFB* in fibroblasts from tumor (red) and adjacent-normal (blue) tissues. **G.** Expression of *NENF*, *HDGF*, and *BMP7* in epithelial cells from tumor and adjacent-normal tissues. These ligands are predicted to mediate epithelial-to-fibroblast signaling.

In adjacent-normal lung tissue, macrophages emerged as a growth factor hub interacting with T cells, endothelial cells, dendritic cells, and charecterized by a strong autocrine signaling loop. Epithelial cells also showed many outgoing growth factor interactions mainly towards endothelial and T cells (Figure 4A). These features are consistent with well-structured barrier tissue maintained by immune surveillance^36^.

In contrast, in the LUSC tumor network macrophages lost their central role. T cells showed an increased autocrine signaling, and CAFs replaced macrophages as key nodes in the network, forming an autocrine signaling loop and establishing interactions with T cells, and cancer epithelial cells. These changes aligned with cold fibrosis, characterized by macrophage depletion and an increase in autocrine-signaling CAFs, similar to the fibrotic scar network architecture observed in myofibroblasts in myocardial infarction^14^. This network architecture is distinct from non-CIAC breast cancer, where fibroblasts lead the signaling hierarchy but have strong interactions with macrophages, indicative of a hot fibrosis state^19^.

To further emphasize tumor-specific signaling, we performed a fold-change-based network analysis comparing tumor (T) and adjacent-normal (N) tissues. To focus on dominant contributors to the LUSC TME, we limited our analysis to cell types that were part of the top 90% strongest outgoing signaling of the total cells in either condition (Figure 4B-C, Methods). In this network, node size represents the fold change in cell type abundance between tumor and adjacent-normal tissue (defined as the percentage of total cells per condition), and edge thickness indicates the fold change in inferred ligand-receptor-mediated communication between cell types.

This network highlighted interactions among T cells, epithelial cells, and CAFs, suggesting that the cold fibrotic stroma is engaged functionally with the hot tumor immune compartment (either to activate or repress immune surveillance)^27,37–39^. CAFs are the dominant signaling hub in the TME, displaying an increase in both incoming and outgoing signaling interactions (∼4-fold increase) (Figure 4C-D). Additionally, T cells showed an >2-fold increase in outgoing signals. Cancer epithelial cells increased only in outgoing signals relative to adjacent-normal epithelium. All outgoing interactions of macrophages were reduced, as expected in the context of a cold fibrosis microenviroment^14^.

These changes suggested the existence of a crosstalk between the fibrotic stroma and adaptive immune cells that occurs in the tumor context. A prominent feature of the LUSC network was the amplification of CAF autocrine signaling in tumors (13.6 fold increase) (Figure 4C), which is also a characteristic of cold fibrosis previously observed in myocardial infarction myofibroblasts^14^ and disease-associated fibroblasts (activated stellate cells) in liver cirrhosis^17^.

To identify potential therapeutic targets within the LUSC TME, we systematically mapped growth factor-mediated interactions in the fold-change based network. Analysis in Figure 4E visualizes growth factor signaling patterns in the LUSC TME, which includes both pleiotropic and exclusive ligands. Among the pleiotropic ligands, *TGFB1* participated in signals between all cell types, including T cells, CAFs, and epithelial cells. In contrast, *TIMP1*, *IHNBA*, and *GMFB* were genes exclusive to CAF in-coming signals, including the CAF autocrine loop, and showed increased gene expression levels in CAFs (Figure 4E-F).

Epithelial-CAF interactions involved heightened levels of *NENF*, *HDGF*, and *BMP7* signaling in tumor epithelial cells (Figure 4E, G). T cell-CAF interactions included factors such as *TGFB1*, *AREG*, *CXCL12* and *IL11*. T cells signal to CAFs via *TGFB1* and *AREG*, and in return CAFs signal through *CXCL12* and *IL11* (Figure 4E). *CXCL12* and *IL11* driven CAF-T cell interactions were distinct from cancer epithelial-CAF interactions, suggesting the existence of additional axes of communication in the LUSC TME. These data point to a set of epithelial, CAF and T cell derived ligands, such as *TIMP1*, *IHNBA*, *GMFB*, *NENF*, *HDGF*, *CXCL12* and *IL11*, whose autocrine and paracrine signaling roles represent potential targets for disrupting the LUSC cold fibrotic TME.

To determine whether the autocrine signaling molecules identified in LUSC CAFs are unique to this tumor type or shared across other CIACs, we examined the expression of *TIMP1*, *INHBA*, *TGFB1*, and *GMFB* in fibroblasts from tumor and normal samples of GAC ^34^and EAC^33^ (Figure S4A). In GAC, *TIMP1* and *TGFB1* were upregulated in tumor-derived CAFs, while *INHBA* and *GMFB* were not detectably expressed in either normal or tumor fibroblasts. In EAC, *TIMP1*, *TGFB1*, and *INHBA* all showed increased expression in tumor-derived fibroblasts relative to their normal counterparts, whereas *GMFB* remained undetected.

Thus *TIMP1* and *TGFB1* represent conserved autocrine signaling molecules across fibroblasts in all three CIACs examined, *INHBA* being selectively shared between LUSC and EAC, while *GMFB* being uniquely associated with CAFs in the LUSC TME. Together, these results identified both shared and tumor-specific components of CAF autocrine signaling in CIACs, pointing to potential mechanisms supporting CAF self-maintenance in the TME of CIACs.

## Discussion

We present a single-cell analysis of LUSC adjacent-normal and tumor samples from 16 treatment-naive patients and analyze its network architecture and cell-task tradeoffs in light of concepts from non-cancer fibrosis. The LUSC TME is enriched with CAFs and has reduced macrophages, resembling cold fibrosis. LUSC CAFs acquire two new functions compared to adjacent-normal fibroblasts: myofibroblast ECM production and immune interaction. The LUSC immune compartment is enriched with T and B cells and shows strong CAF-T cell growth factor signaling interactions. The LUSC TME circuit differs from the previously characterized non-CIAC breast cancer TME circuit (from multiple breast cancer subtypes) which resembles hot fibrosis with key roles for macrophage-fibroblast interactions^19^.

The concept of cold fibrosis in the TME enhances the classical histological designations of ECM-rich tumor regions as fibro-inflammatory or desmoplastic^7,8^. Cold fibrosis refers to a specific cellular composition and cell-circuit architecture, characterized by macrophage depletion and myofibroblast (or CAF) expansion sustained by autocrine growth factor signaling^12,14^. Cold fibrosis has been validated *in vivo* by our group and others, in settings of myocardial infarction^14,15^ and human kidney transplantation^14,15^. Here, we propose that ECM-rich regions within solid tumors depleted from macrophages, including those labeled as desmoplastic, can also be described as cold fibrosis, expanding the paradigm of the TME as a chronic wound undergoing repair and regeneration ^40^.

In LUSC, CAFs are significantly expanded and transcriptionally reprogrammed, while macrophages are markedly depleted. CAFs adopt two new cellular states not observed in adjacent-normal lung samples: (1) a contractile, ECM-remodeling myofibroblast program, and (2) a cellular-stress, immune-interacting phenotype. These states mirror the fibroblast phenotypes observed in cold fibrotic scars in the heart, where fibroblasts acquire persistent fibrotic ECM maintenance properties^14^. We find this gene expression program in EAC and GAC, two additional CIACs. Each of the three CIACs showed depletion of quiescent fibroblasts and enrichment of stress-adapted and myofibroblast states. These findings suggest that CIACs may share a core cold fibrosis program.

Our network analysis identifed CAFs as dominant signaling hubs within the LUSC TME. A previous study that analyzed non-CIAC breast cancer subtypes, including: estrogen receptor-positive (ER^+^), HER2^+^, and triple-negative breast cancer (TNBC), demonstrated that fibroblasts also function as key signaling hubs, but engage in strong paracrine circuits with tumor associated macrophages, consistent with a hot fibrosis state^19^. In LUSC, CAFs operate independently of macrophages, relying instead on autocrine loops and crosstalk with T cells and cancer epithelial cells. This topology is similar to the myofibroblast-centric signaling architecture seen in cold cardiac (non cancerous) fibrosis^14^. It would be interesting to further investigate whether different CIACs exhibit conserved or distinct patterns of cell-cell interactions.

In heart and liver fibrosis, the myofibroblast autocrine loop was a target for preventing or reducing fibrosis in mice^14,17^. We therefore explored the autocrine loop of CAFs in LUSC. We identified several fibroblast-derived autocrine growth factors including *TIMP1*, *INHBA*, *TGFB1*, and *GMFB*. TIMP1 has been shown to promote proliferation of scleroderma fibroblasts and hepatic stellate cells^41,42^. Its capacity to stimulate proliferation was observed independent of its canonical role as an MMP-inhibitor, as demonstrated in keratinocytes^43,44^. We previously demonstrated that TIMP1 acts as a cardiac myofibroblast growth factor in fibrosis, where it supports myofibroblast proliferation through its N-terminal domain^14,45^. TIMP1 neutralizing antibodies reduced fibrosis after heart injury in mice^14^.

Other CAF autocrine factors in LUSC include *INHBA*, which encodes Activin A, and drives fibroblast proliferation and ECM deposition in lung fibrosis and cancer ^46–48^. *TGFB1* acts in both autocrine and paracrine manners, and is well known for its dual role in matrix production and T cell suppression^49–52^ - although its pleiotropic nature suggests that targeting it is challenging. GMFB, while previously studied in glial inflammation and hepatocellular carcinoma^53,54^, appears here as a LUSC-selective autocrine ligand of CAFs.

Some of these autocrine factors are also elevated in the TME of other CIACs. *TIMP1* and *TGFB1* were upregulated in both EAC and GAC CAFs; *INHBA* was shared between LUSC and EAC, while *GMFB* was unique to LUSC. Together, these factors support a self-reinforcing CAF loop that may be targetable in LUSC and other CIACs, just as in the fibrotic heart and liver^14,17^.

One question for further research is the spatial organization of cold fibrosis in the TME. Spatial proteomics has shown that CAFs in LUSC form barriers around tumor nests that exclude T cells^22^. Our data suggests that the LUSC TME contains an adaptive immune infiltrate, with increased T and B cell numbers relative to the adjacent-normal tissue. This “hot” immune environment coexists with cold fibrosis, supporting a model where spatially and functionally separated immune and fibrotic compartments define the TME. Our network analysis reveals reciprocal signaling that may support this spatial segregation. T cells signal to CAFs via *TGFB1* and *AREG*, both of which can induce fibroblast activation and profibrotic gene programs^49,55^. CAFs in turn secrete *CXCL12*, a chemokine previously shown to restrict T cell infiltration into tumors, and *IL11*, a cytokine that promotes profibrotic signaling^56,57^. These interactions may be consistent with an avoidance circuit in which immune and stromal compartments remain functionally engaged but spatially distinct.

Previous attempts to modulate the fibrotic stroma in cancer focused on mitigating the effects of CAFs and other fibroblasts, such as their ECM and signaling functions. In pancreatic ductal adenocarcinoma (PDAC), inhibition of lysyl oxidase (LOX) enzymes, key mediators of collagen cross-linking, reduced ECM abundance, enhanced chemotherapy response, and increased survival in mouse models^58^. In the liver metastatic niche of PDAC, it was shown that macrophage-driven activation of hepatic stellate cells (HSCs) supports metastatic cell proliferation. Blocking this circuit by reducing macrophage numbers, blocking HSC activation or by inhibiting HSC-derived POSTN, limited metastatic outgrowth^59^. In glioblastoma, treatment with surgical resection, ionizing radiation therapy, or CSF1R inhibition lead to fibrotic scar formation, which provides a niche for dormant tumor cells to survive and later drive recurrence in adjacent healthy tissue^60^. Combined inhibition of TGFB1, CSF1R, together with an anti-inflammatory agent reduced fibrotic scars and improved survival in preclinical animal models ^60^.

In light of our findings, we propose an alternative strategy: targeting the autocrine loop of cold-fibrosis-like CAFs to drive deletion of this cell population, rather than merely suppressing ECM deposition or CAF phenotypes. This approach could eliminate the full spectrum of pro-fibrotic and immune-modulatory signals secreted by CAFs in a single step. Importantly, the effects of CAF depletion may depend on tumor context; for example, in pancreatic cancer, genetic ablation of αSMA⁺ myofibroblasts led to more aggressive, immune-suppressed tumors and reduced survival^61^. These findings highlight the need to carefully evaluate the consequences of CAF targeting in each specific tumor setting. Framing the LUSC TME as an example case of cold fibrosis not only provides a conceptual framework but also reveals testable vulnerabilities for therapeutic interventions (cold fibrosis relies on the autocrine loop to support the fibroblasts). In summary, we describe a cold fibrotic architecture in the LUSC TME, defined by autocrine self-maintenance of activated CAFs, and concomitant loss of macrophage support. This fibrotic program coexists with a hot adaptive immune infiltrate and is found in other CIACs, such as EAC and a subset of GAC. The parallels between LUSC, fibrotic heart scars, and other CIACs suggest that cold fibrosis may represent a conserved cell-circuit in chronic tissue remodeling. By identifying CAF-specific autocrine growth factors and unveiling shared mechanisms across disease contexts, we provide a framework for future therapeutic strategies aimed at collapsing fibrotic structures within tumors.

## Methods

### Human lung squamous cell carcinoma and adjacent-normal tissue samples

Fresh tissue specimens were obtained from patients who provided written, informed consent and were diagnosed with treatment-naive lung squamous cell carcinoma (Research Institute - McGill University Health Centre REB # 2014-1119) at the time of surgical resection. The samples encompassed visually identified malignant tumors and distal apparent-normal regions from material assessed by a trained clinical pathologist directly following resection. Regions from which samples were taken were inked on the remaining fresh gross specimen at the time of collection. Histological diagnosis was performed on H&E-stained sections of formalin-fixed paraffin-embedded tissue blocks following the identification of the previously inked regions and corroborated by the consensus of two expert pathologists. Only tissue specimens with confirmed histological diagnosis were used in subsequent analyses. Collected tissue specimens were placed in a cold medium (RPMI (Invitrogen) supplemented with Primocin (InvivoGen) and gentamycin (Invitrogen) for single-cell RNA sequence processing.

### Single-cell RNAseq Methods Single-cell dissociation

Tissue specimens were dissected to remove necrotic areas, minced, and digested in 5 mL of Advanced DMEM/F12 containing 10 mg Collagenase Type 3 (Worthington) and 500U Hyaluronidase (Sigma) in a C-tube (Miltenyi) using the gentleMACS Octo Dissociator (Miltenyi). The single-cell suspension was resuspended in PBS and 1mM DTT, strained through a 100um cell strainer (Fisher), and spun down (500xg, 5 minutes, 4°C). Cells were resuspended in 0.25% Trypsin-EDTA (Invitrogen) and incubated for 5 minutes at 37°C, followed by the addition of 10% fetal bovine serum to inactivate trypsin. The cell pellet (500xg, 5 minutes, 4°C) was resuspended in 2.5U Dispase/10ug DNAse buffer and incubated for 5 minutes at 37°C. The buffer was inactivated by adding excess PBS and the homogenate was strained (40um, Fisher) prior to centrifugation (500xg, 5 minutes, 4°C). Red blood cells were lysed using ACK Lysing Buffer (Gibco) for 5 minutes at room temperature, followed by the addition of excess PBS, prior to centrifugation (500xg, 5 minutes, 4°C). The cell pellet was finally washed twice with 2% fetal bovine serum in PBS prior to proceeding with single-cell capture on the 10x Genomics platform.

### Single-cell suspension quality assessment

Before credentialing the cell suspension, the cells were filtered through a 40um FLOWMI cell strainer (SP Bel-Art; H13680-0040). Whenever necessary, centrifugation of the cells was carried out at 300xg for 11 minutes. Single-cell viability and the presence of debris and erythrocytes in the single-cell suspension were assessed prior to single-cell capturing. Upon adequate viability (i.e. lack of debris and erythrocytes), cells were captured on the 10x Genomics platform.

Cell viability was tested using the “LIVE/DEAD Viability/Cytotoxicity Kit for mammalian cells” that contained Ethidium Homodimer-1 and Calcein-AM stain (ThermoFisher; L-3224) dyes. First, a viability stain mix was reconstituted by mixing 0.5 ul of 4mM Calcein-AM, 2ul of 2mM Ethidium Homodimer-1, and 100ul of PBS. A 5ul of cell suspension was then resuspended in 5ul of viability stain mix and the solution was incubated at room temperature for 10 minutes. The sample viability was verified using a hemocytometer (INCYTO C-Chip; DHC-N01-5) through GFP (for the Calcein-AM) and RFP (for the Ethidium Homodimer-1) channels on an EVOS FL Auto Fluorescent microscope (ThermoFisher). Viability was expressed as the percentage of live cells (Calcein-AM / GFP positive cells) over the sum of live (Calcein-AM / GFP positive cells) and dead cells (Ethidium Homodimer-1 / RFP positive) cells.

Erythrocyte contamination was assessed by staining the cells with the cell-permeable DNA dye DRAQ5 (ThermoFisher; 65-0880-92). A nuclear staining mix was made by diluting the DRAQ5 stock solution (5mM) down to 5uM with 1x PBS. Afterwards, 5ul of cell suspension was re-suspended in 5ul of nuclear stain mix and the solution was incubated at room temperature for 5 minutes. The nuclear stain was visualized using a hemocytometer (INCYTO C-Chip; DHC-N01-5) through the Cy5 channel on an EVOS FL Auto Fluorescent microscope (ThermoFisher). Erythrocyte contamination was expressed as the percentage of “round donut-shaped DRAQ5 negative objects on bright-field” over the sum of “round donut-shaped DRAQ5 negative objects on bright-field” and “nuclear stained DRAQ5 positive cells”.

Additionally, we assessed the cell suspension for the presence of any other contaminants/debris as well as contaminants that might interfere with capture on the microfluidic chip such as large debris. The percentage of debris present in the sample was expressed as follows: Percentage of “observed non-cell objects on bright-field” over the sum of “observed objects on bright-field” and “observed cells marked on the fluorescent channels”. A sample was deemed adequate for capturing if “cell viability” was ≥ 70%, “erythrocyte contamination” was ≤ 10%, and “debris percentage” was ≤ 30%. The cell concentration in the cell suspension, measured by counting the number of “Calcein-AM / GFP positive cells” and “Ethidium Homodimer-1 / RFP positive cells” in the large 4 squares on each corner of the hemocytometer, was calculated as follows: The number of cells/ul =[(Calcein-AM / GFP positive cells + Ethidium Homodimer-1 / RFP positive cells) / 4] * 10 * 2 where 10 was the dilution factor on the hematocytometer and 2 was the dilution factor when the cell suspension was mixed with the dye solution.

### Single-cell capturing

Single cells were captured on the 10x Genomics platform. Single-cell 3’ end gene expression profiling was carried out according to the “Chromium Next GEM Single Cell 3’ Reagent Kits v3.1” protocol and recommended reagents. Single-cell Copy Number Variation was queried using the “Chromium Single Cell DNA Reagent Kits” protocol and recommended reagents. Of note the CNV kit described above is currently discontinued. The sequencing libraries were created as per the above protocols with the modifications presented in the following section.

### Sequencing of the 10x single-cell libraries

Libraries were quantified using a LightCycler 480 Real-Time PCR instrument (Roche) and the KAPA library quantification kit (Roche) with triplicate measurements. Library quantification values were used both for the MGI library conversion and for Illumina sequencing normalization.

Libraries sequenced on MGI (MGI Tech) were converted after 10x library construction to be compatible with MGI sequencers using the MGIEasy Universal Library Conversion Kit. The kit circularizes the libraries making them compatible with MGI systems. To sequence the circularized libraries, they were first amplified by rolling circle amplification, resulting in a long DNA strand that individually folds into a tight ball (i.e. a DNA nanoball) where one library fragment results in one DNA nanoball. Before loading into the flowcells, the amplified nanoballs were quantified with a Qubit ssDNA HS Assay kit (ThermoFisher), normalized, and loaded onto the sequencing flowcell using the auto-loader method (auto-loader MGI-DL-200R). The flowcells have a functionalized surface that captures and immobilizes the nanoballs in a grid pattern. Typically, two libraries were loaded per lane for the single-cell RNA libraries. The DNBSEQ-G400RS PE100 MGI kit with App-A primers was used for single-cell RNA library sequencing. The DNBSEQ-G400RS PE150 MGI kit with App-A primers was used for single-cell DNA library sequencing.

The flowcells were sequenced on a DNBSEQ-G400 MGI sequencer. Single-cell RNA libraries were sequenced as follows: 28 cycles for read1, 150 cycles for read2, and 8 cycles for the i7 index. Single-cell DNA libraries were sequenced as follows: 151 cycles for Read1, 151 cycles for Read2, and 8 cycles for the i5 index. Because libraries must be color-balanced for all cycles sequenced in order to maintain a minimum ratio of 0.125 for each base at each cycle, color-balanced single index adapters (10x Genomics) were used for libraries sequenced on MGI.

A subset of 2 libraries was sequenced on the Illumina NovaSeq 6000 platform using S4 flowcells after splitting the original cDNA. To ensure uniform loading of the libraries, a preliminary pool was sequenced on Illumina iSeq, and the library proportions were readjusted accordingly. Although the MGI sequencer has onboard capability to demultiplex samples, we chose to use independent tools to demultiplex the raw fastq files for each lane to give us the flexibility to reprocess if needed. The fastq files generated using the balanced single index adapters were merged for each library after demultiplexing. The MGI runs were mainly demultiplexed by fastq-multx^62^ (https://github.com/brwnj/fastq-multx) but also using fgbio/DemuxFastqs (http://fulcrumgenomics.github.io/fgbio/tools/latest/DemuxFastqs.html). In both instances, we used a mismatch of 1. Illumina runs were demultiplexed using the standard bcl2fastq tool.

### Read processing and alignment

After polyA-trimming via cutadapt (v3.2)^63^, reads were pseudo-aligned to the GRCh38 reference transcriptome (ENSEMBL release 96) with kallisto (v0.46.2)^63,64^ using the default kmer size of 31. The pseudo-aligned reads were processed into a cell-by-gene count matrix using bustools (0.40.0)^65^. Cell barcodes were filtered using the whitelist (v3) provided by 10xGenomics. All further processing was done in scanpy (v.1.7.1)^65,66^.

### Quality control and normalization

Quality control was performed for each sample independently as follows. Cell barcodes with less than 1000 counts or less than 500 genes expressed or with more than 10% mitochondrial gene expression were removed. Doublet cells were identified using scrublet^67^, and any cell barcode with a scrublet score > 0.2 was removed. Only coding genes were retained in the final count matrix. Expression profiles were normalized by total counts, the 4000 most highly variable genes being identified^68^, renormalized, log-transformed, and z-scored. The data were projected onto the first 50 principal components. This resulted in a data set with 197,232 cells by 35,606 genes. To remove any batch effects, the Harmony algorithm^68,69^ was applied on the first 50 principal components with a maximum of 25 iterations. A nearest neighbor graph (k=15) was calculated on the Harmony-corrected principal components space. Datasets were visualized in 2D via UMAP93 and initialized with PAGA^70^ coordinates. The nearest neighbor graph was clustered with the Leiden algorithm^70,71^.

Single-cell RNA-sequencing data were imported from an annotated .h5ad file using Seurat (v4.3.0)^72,73^ and associated packages. The original object was converted to Seurat format using the *Convert* function from SeuratDisk, and subsequently loaded into R as a Seurat object with *LoadH5Seurat* function. The processed Seurat object was saved as an .rds file for downstream analyses.

### Cell type classification

Cell type annotation was performed using a hybrid approach that combined unsupervised clustering-derived marker gene identification with manual curation based on known cell type-specific markers. First, differentially expressed genes for each cluster were identified using the *FindAllMarkers* function from the Seurat R package^72,73^. The analysis was restricted to genes with positive log fold change, a minimum log_2_ fold change of 0.25 (logfc.threshold = 0.25), and a minimum expression threshold of 25% of cells within a cluster (min.pct = 0.25). The resulting cluster-specific markers (Table S1) were exported for downstream inspection. These markers were then compared to established cell-type specific gene signatures to assign biological identities to each cluster.

### Cell type-specific differential gene expression analysis and visualization

Differentially expressed genes were analyzed separately for each annotated cell type using a custom R workflow based on the Seurat v4 package^72,73^. A Seurat object containing single-cell RNA-seq data from both tumor (“T”) and adjacent-normal (“N”) samples was used as input. For each cell type, a subset Seurat object was created, and pairwise differential expression analysis between tumor and adjacent-normal cells was performed using the Wilcoxon rank-sum test implemented in the *FindMarkers* function (test.use = “wilcox”). Only genes expressed in at least 30% of cells (min.pct = 0.3) and with an absolute log_2_ fold change greater than 0.4 were considered. P-values were adjusted using the Benjamini-Hochberg method for multiple testing correction.

The output for each cell type was stored as an element in a list, and results were merged into a single data frame. Genes were classified into three categories based on statistical significance: Upregulated (adjusted p-value < 0.01 and log_2_ fold change > 1), Downregulated (adjusted p-value < 0.01 and log_2_ fold change < -1), or Not Significant.

To visualize the number of differentially expressed genes per cell type, up- and down-regulated gene counts were computed for each subset and plotted as a scatter plot with log_10_ scaling on both axes. Volcano plots were then generated for each cell type to visualize the relationship between fold change and statistical significance. These plots displayed log_2_ fold change on the x-axis and -log_10_ adjusted p-value on the y-axis, with dashed lines indicating the thresholds for significance (log_2_FC = ±1, p_adj = 0.01).

### Fibroblast subclustering and archetype scoring

The fibroblast cluster was subsetted using Seurat^72,73^. The resulting subset was normalized using *NormalizeData*, followed by identification of variable genes with *FindVariableFeatures*, scaling with *ScaleData*, and dimensionality reduction using *RunPCA* function. Elbow plot analysis was performed using the *ElbowPlot* function to guide dimensionality selection, and the first 24 principal components (PCs), which captured at least 80% of the variance, were used for clustering and visualization. Clustering was performed using *FindNeighbors*(dims = 1:24) and *FindClusters*(resolution = 0.5). UMAP embeddings were generated with *RunUMAP* function (dims = 1:24). To investigate fibroblast functional diversity, we defined four archetype-specific gene signatures (Arch1-4) based on genes enriched in specific ParTI-defined Pareto archetypes (median difference > 1, p < 0.05) (Figure 3). The number of genes included in each signature was: Archetype #1 = 98 genes, archetype #2 = 23 genes, archetype #3 = 108 genes, and archetype #4 = 42 genes (Table S6). Each gene set was scored individually using Seurat’s *AddModuleScore* function (nbin = 12), and visualized using *FeaturePlot* on the fibroblast UMAP.

### Epithelial cells subclustering and archetype scoring

Epithelial cells cluster was subsetted using Seurat^72,73^. Subsequent analyses were performed in R using the Seurat package. The subset object was normalized using the *NormalizeData* function, followed by identification of variable features with *FindVariableFeatures* and scaling with *ScaleData*. PCA analysis was performed to reduce dimensionality with *RunPCA*, and the variance explained by each PC was calculated to determine the number of PCs required to capture 80% of the variance. An elbow plot was generated to mark the PC cutoff for 80% cumulative variance. Based on this analysis, the first 19 PCs were selected for downstream clustering and visualization. Cells were reclustered using *FindNeighbors* and *FindClusters* functions at a resolution of 0.1, and visualized via UMAP with the *RunUMAP* function. UMAP plots were generated to show epithelial cell clusters and distribution by diagnosis. Marker genes for each cluster were identified using *FindAllMarkers* function with thresholds of log_2_ fold-change > 0.25 and minimum expression in 25% of cells (min.pct = 0.25). The top three marker genes per cluster were extracted and visualized using dot plots (*DotPlot* function). To investigate epithelial cells functional diversity, we defined four archetype-specific gene signatures (1-4) based on genes enriched in specific ParTI-defined Pareto archetypes (median difference > 1.5, p < 0.01) (Figure 2, Table S4). Each gene set was scored individually using Seurat’s *AddModuleScore* function (nbin = 12), and visualized using *FeaturePlot* function on the fibroblast UMAP.

### Pareto analysis

We analyzed the epithelial cells and fibroblasts scRNAseq data and performed Pareto Optimality analysis to infer the trade-offs and tasks that the cells specialize in. The raw data includes 29,570 genes for epithelial cells and 25,535 genes for fibroblasts. After downsampling, we included in the analysis 10,000 epithelial cells (5156 from adjacent-normal tissue and 4844 from tumor tissue) and 5229 fibroblasts (1290 from adjacent-normal tissue and 3939 from tumor tissue). We used Sanity^74^, a method to normalize scRNAseq data and to infer the transcription activity of the genes. Sanity is a Bayesian procedure for normalizing scRNAseq data from first principles. Following the Sanity normalization, genes with low mean expression and low variation were removed. Only genes with log10 mean expression larger than the median expression and standard deviation in the top 25th percentile were considered. We ended up with 6618 genes for the epithelial cells and 5246 for the fibroblasts. We next used the ParTI package in Matlab to fit the data of each cell type population from each condition to a polytope. We find that epithelial cells are distributed within a 4-vertex simplex (p-value <10^−5^) and fibroblasts also best fit with a tradeoff among 4 tasks (p-value = 0.03). We next used ParTI to find enriched genes for all the archetypes to infer the tasks the cells specialize in. We quantified the distance between each cell and each archetype in PCA space using the Euclidean norm. For each archetype, we divided the distance range-from the closest to the farthest cell-into five equal-width bins. Cells were assigned to bins based on their distance to the archetype, and the proportion of tumor versus adjacent-normal cells within each bin was computed.

### NicheNet analysis

To investigate intercellular communication across annotated cell populations, we applied the NicheNet framework^35,75,76^, which integrates prior knowledge of ligand-receptor interactions and downstream target gene regulation. A Seurat object containing normalized expression data and cell-type annotations was used as input. For each receiver cell population (e.g., macrophages, fibroblasts), genes of interest (GOIs) were defined based on differential expression (log₂ fold change ≥ 0.25, adjusted p-value ≤ 0.05, and expression in ≥10% of cells). These GOIs were then matched against the NicheNet ligand-target matrix to infer candidate ligands from sender cell populations that could explain observed gene expression changes. Pairwise analyses were performed across all sender-receiver cluster combinations to identify ligands with strong predicted regulatory activity, as quantified by Pearson correlation between the predicted and observed GOI expression. Ligands ranking in the top 20th percentile were considered active and were further analyzed for expression patterns, regulatory potential, and receptor availability in receiver populations.

### Cell-cell communication network analysis

The pairwise interaction between cell types (macrophages, fibroblasts, etc.) was calculated independently for each sample (adjacent-normal and tumor). Genes of interest (GOIs) were selected from a list of growth factors (GO:0008083). The interaction score was calculated as the Pearson coefficient calculated by the ‘predict_ligand_activities’ function, using the GOIs of the genes that were expressed in the ‘receiver’ clusters as targets.

To analyze the interaction strength (weighted interaction) between the various cell types, we first estimated the percentage of every cell type from the total in each sample. Cell types that contributed less than 1% to the total cell count were ignored. These percentages were then used to determine the sizes of the vertices in the graph. Next, the edge weights were calculated by multiplying three terms: (1) % of the sender cell (%𝐶𝑆) - the number of “broadcasting antennas”), (2) the regulatory potential of the sent ligands as estimated from NicheNet (summed over all ligand-receptors pairs: ∑𝑙∈{𝐿𝑅𝑝𝑎𝑖𝑟𝑠}𝜚_S_→𝑅), and (3) % of the receiver cell (the number of “listening antennas”: 𝐶𝑅:

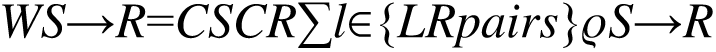

In the fold change network of the tumor-gained functions, vertex sizes were calculated as the fold change in cell % in tumor vs adjacent-normal and edge weights as the fold change between an edge in the tumor network and its corresponding edge in the adjacent-normal network. In the fold change network of the tumor lost functions the inverse ratios were used. All calculations and visualizations were performed using custom scripts in Mathematica 14.0.

### Statistical analyses

Specific details of statistical tests used for each figure panel can be found in the appropriate figure legend. Statistical analysis of paired tumor vs adjacent-normal cell type abundance (%; Figure 1E), was performed calculated by using a paired two-tailed Student’s t-test using Prism v10.4. Paired comparisons of fibroblast archetype module scores between tumor and adjacent-normal samples (Figure 3G, H) were assessed using the two-sided paired Wilcoxon signed-rank test, implemented in R using the *wilcox.test* function. Individual data points represent data derived from different biological repeats (unless mentioned differently in figure legends). Statistical analysis is derived from the biological repeats of an experiment. Measurements are reported as the mean and the error bars denote the S.E.M. throughout the study (unless mentioned differently in figure legends). p-values are presented on figures in the appropriate comparison location. Statistical analysis of Pareto archetype inference and enrichment of genes was done using the ParTI package ^18^ in Matlab (R2022a).

## Supporting information

Single-cell mRNA-sequencing, cell type marker genes, related to Figure 1

Single-cell mRNA-sequencing, cell type specific differentially expressed genes of tumor vs adjacent-normal cells, related to Figure 1

Single-cell mRNA-sequencing, epithelial cells Seurat cluster defining genes, related to Figure 2

ParTi analysis of enriched archetype genes near epithelial cells vertices (tumor and adjacent-normal cells), related to Figure 2

ParTi analysis of enriched archetype genes near fibroblast vertices (tumor and adjacent-normal cells), related to Figure 3

Single-cell mRNA-sequencing, epithelial cells Seurat cluster defining genes, related to Figure 3

## Acknowledgments

This study was supported by Grand Challenge Cancer Research UK awards A27145 to TDT, A29077 to UA, A29068 to SH and A29071/A29078 to LF, NCI Outstanding Investigator Award R35 CA197694 to TDT and NCI R50 CA211543 to PG. MA is supported by the Israel Science Foundation (ISF, Grant No. 3372/24); the United States–Israel Binational Science Foundation (BSF, Grant No. 2023231); and is the recipient of the Alon Fellowship, Israel Council for Higher Education.

## Author contributions

Conceptualization: SM, AM, PG, TT, UA; Methodology: SM, AM, SF, MS, MA; Investigation: SM, SF, AM, PG, MS, DG, EW, IBS, YS, DP, JAC, VS, NB, JB, SCB, JR, SO, SMa, MA; Visualization: SM, AM, SF; Funding acquisition: ET, TT, UA; Project administration: PG, TT, UA; Supervision: SH, LF, TT, ET, RSS, UA; Writing - original draft: SM, SF, AM, ET, RSS, UA; Writing - review & editing: All authors.

## Data and materials availability

All data supporting the findings of this study are available within the main text and supplementary materials. Processed single-cell RNA-seq data, including count matrices and metadata, will be deposited in the Gene Expression Omnibus (GEO) and made publicly available upon publication. All custom analysis code will be made available on the AlonLabWIS GitHub repository.

## Supplemental information

**Table S1** Single-cell mRNA-sequencing, cell type marker genes, related to Figure 1

**Table S2** Single-cell mRNA-sequencing, cell type specific differentially expressed genes of tumor vs adjacent-normal cells, related to Figure 1

**Table S3** Single-cell mRNA-sequencing, epithelial cells Seurat cluster defining genes, related to Figure 2

**Table S4** ParTi analysis of enriched archetype genes near epithelial cells vertices (tumor and adjacent-normal cells), related to Figure 2

**Table S5** ParTi analysis of enriched archetype genes near fibroblast vertices (tumor and adjacent-normal cells), related to Figure 3

**Table S6** Single-cell mRNA-sequencing, epithelial cells Seurat cluster defining genes, related to Figure 3

## Figure legends

**Figure S1.**
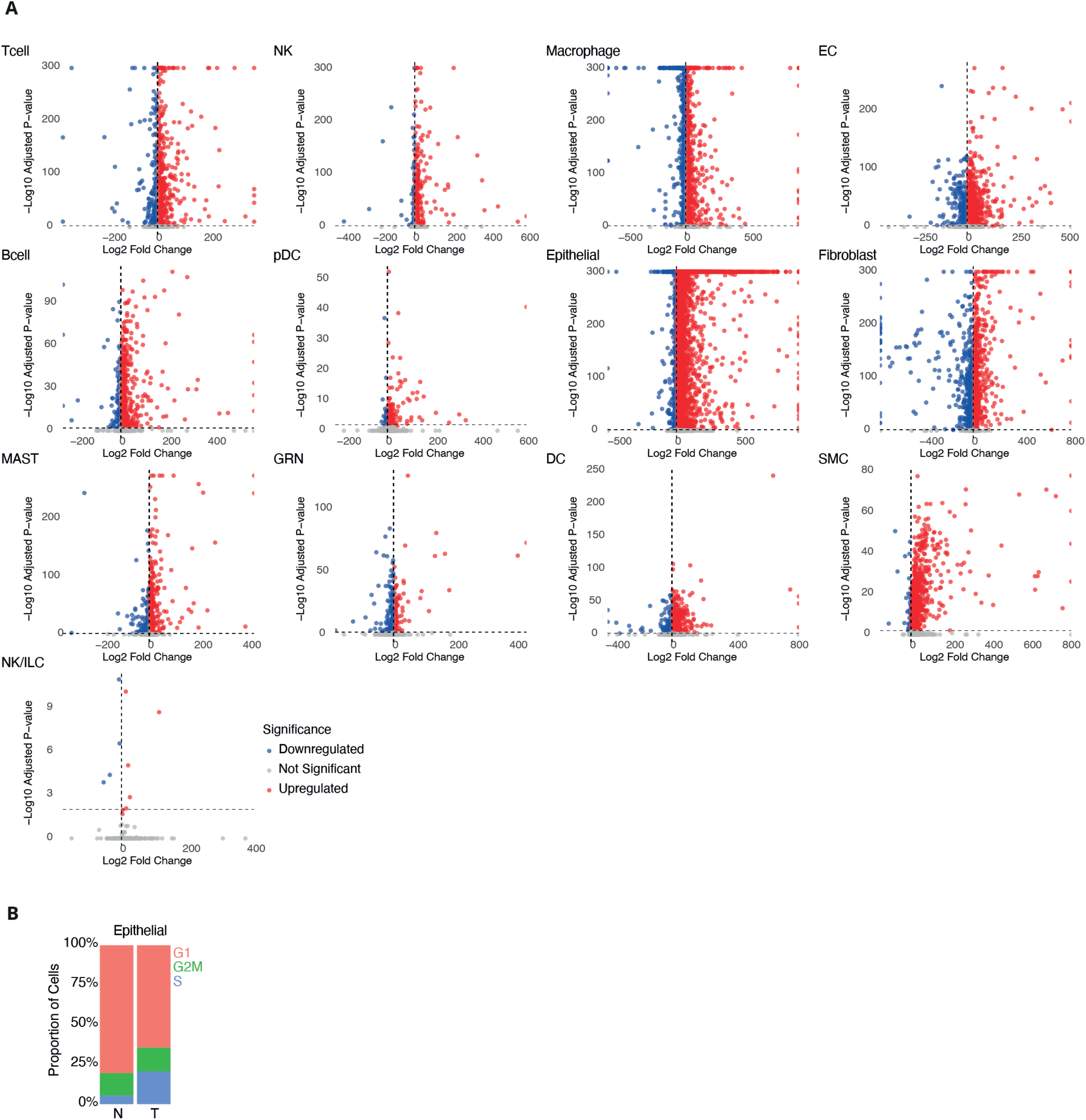
Volcano plots of differentially expressed genes across cell types (tumor vs adjacent-normal), related to Figure 1. **A.** Volcano plots display log2 fold change (x-axis) versus -log10 adjusted p-value (y-axis) for each gene, computed separately for each annotated cell type. Genes were classified as upregulated (adjusted p-value < 0.01 and log2FC > 1, red), downregulated (adjusted p-value < 0.01 and log2FC < -1, blue), or not significant (gray). Dashed vertical lines at log2FC ±1 and a horizontal line at -log10(p_adj) = 2 (equivalent to p_adj = 0.01) indicate significance thresholds used to define the differential expression. Each panel corresponds to one cell type from the dataset. **B.** Relative cell cycle phase distribution in epithelial cells by diagnosis (Methods). Proportions of G1, S, and G2/M phase cells were computed per diagnostic group and displayed as normalized stacked bar plots.

**Figure S2.**
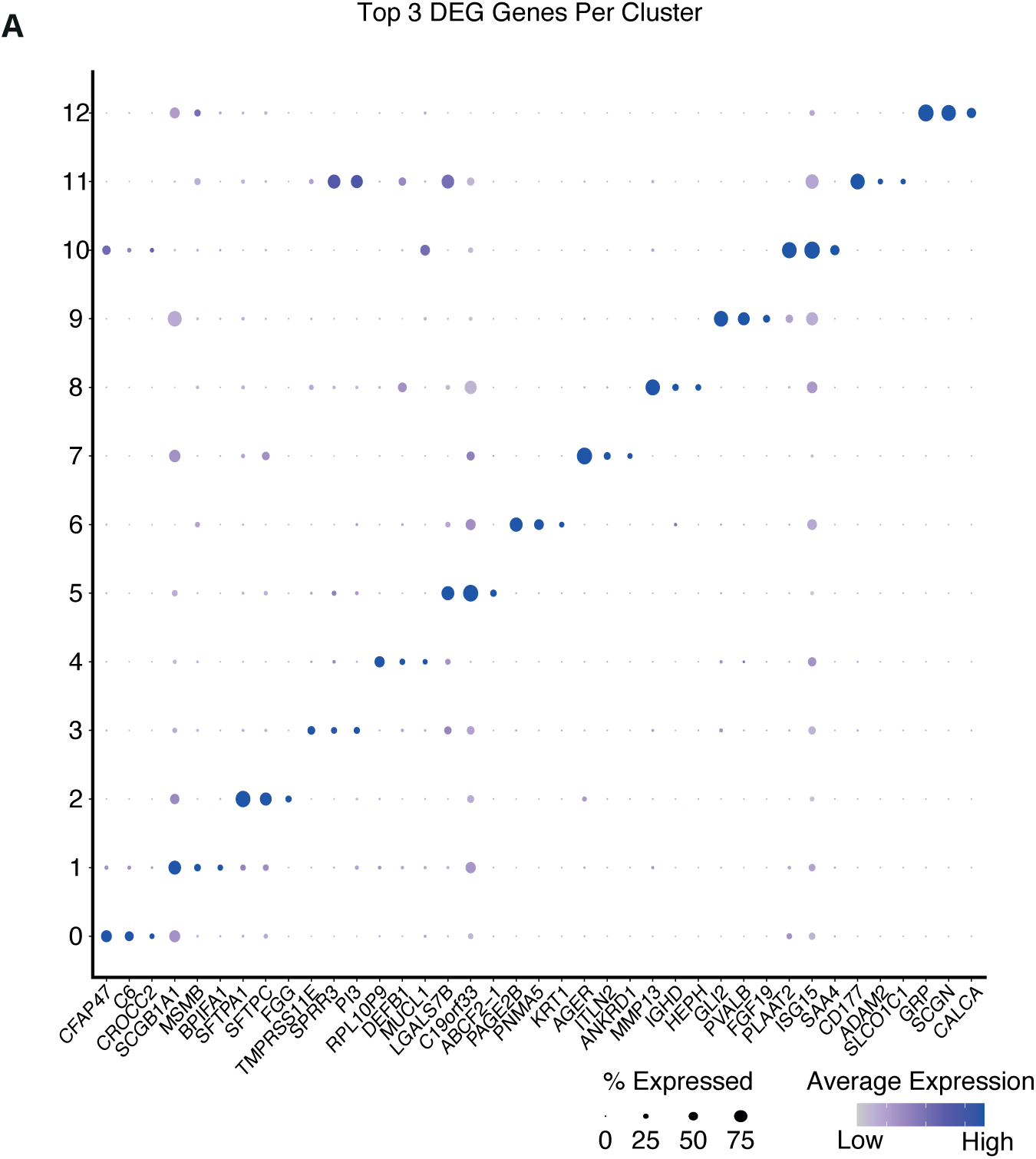
Epithelial cells, Seurat clustering, and Archetype score, related to Figure 2. **A.** Dot plot displaying the top 3 marker genes per epithelial cluster, as identified by differential gene expression analysis (Methods, Table S3). Marker expression patterns highlight transcriptional differences across epithelial subpopulations. Color represents average expression levels per cluster on a linear scale. The circle size represents the percent (%) expressing cells per cluster.

**Figure S3.**
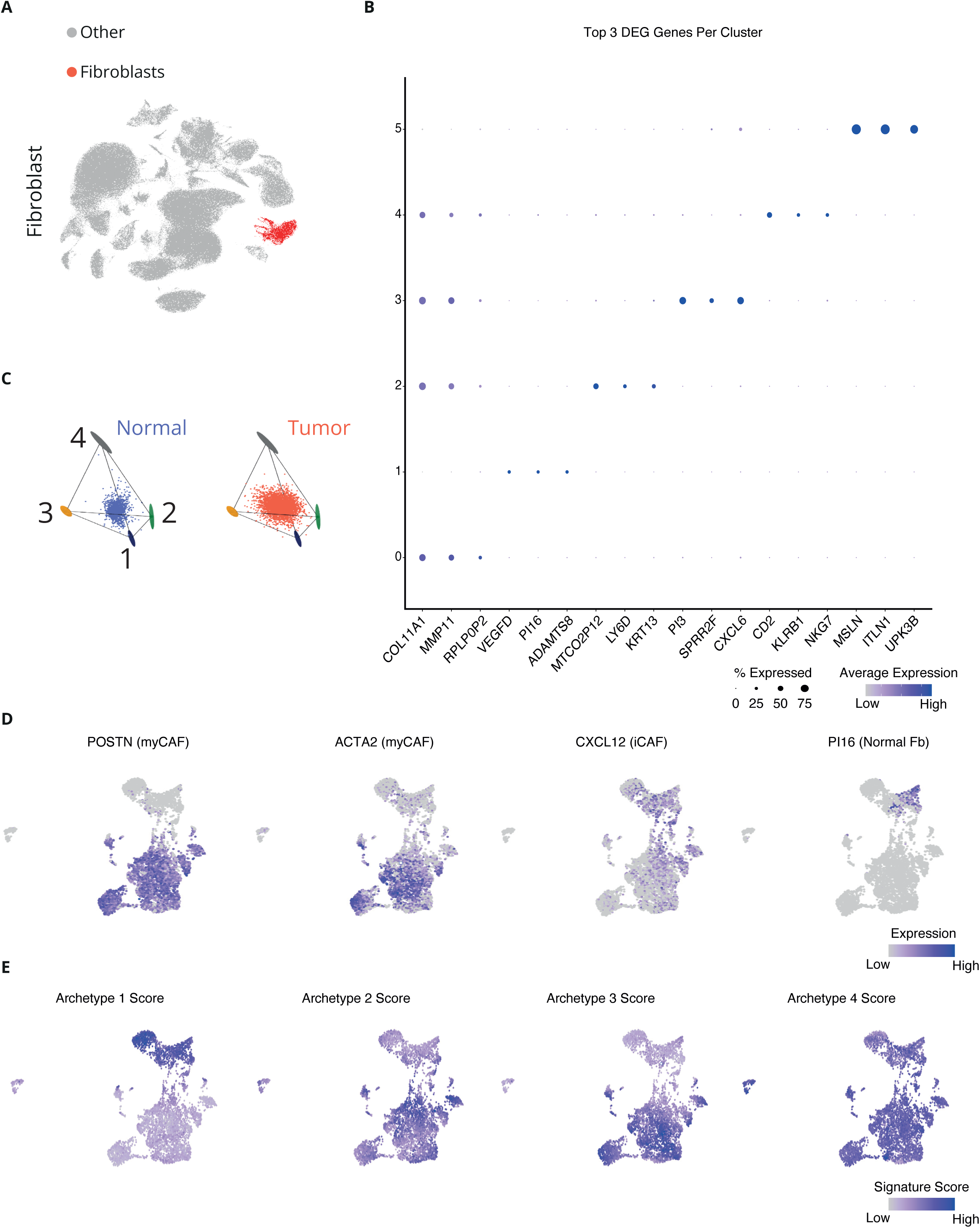
LUSC fibroblast archetypes correspond to a myofibroblast-CAF (myCAF) phenotype, related to Figure 3. **A.** UMAP embedding of all cells, with fibroblasts highlighted in red and all other cell types shown in gray. **B.** Dot plot showing the top 3 marker genes per fibroblast cluster identified by differential gene expression analysis (Methods, Table S5). Color represents average expression per cluster (linear scale); dot size represents the percent (%) of cells expressing the gene. **C.** Feature plots showing the expression of canonical myCAF, inflammatory-CAF (iCAF), and adjacent-normal fibroblasts marker genes: *POSTN*, *ACTA2*, *CXCL12*, *PI16*, respectively. UMAP projections illustrate the distribution of marker expression levels. Color represents expression levels (linear scale). **D.** Module scoring of four fibroblast archetypes based on gene signatures identified through ParTI archetype analysis (Figure 3C, Table S6, Methods). Feature plots show UMAP projections of module scores for each archetype. Cells are colored by module score expression (linear scale), revealing spatial enrichment of distinct fibroblast states.

**Figure S4.**
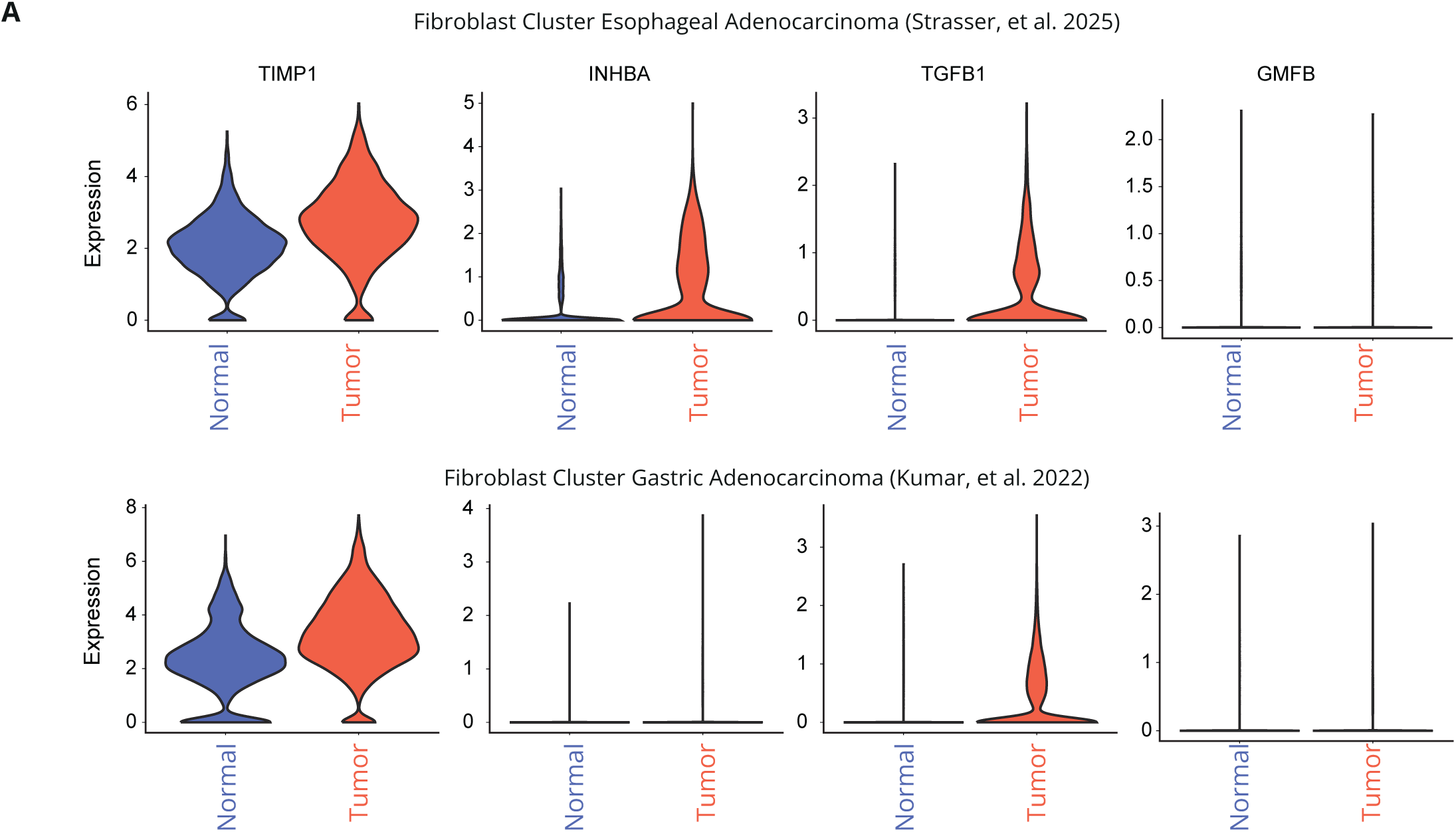
Tumor-enriched expression of CAF autocrine factors in CIACs (EAC and GAC), related to Figure 4. **A**. Expression of *TIMP1*, *INHBA*, *TGFB1*, and *GMFB* in fibroblasts from adjacent-normal and tumor samples in two additional chronic inflammation-associated cancers (CIACs): esophageal adenocarcinoma^33^ (EAC, top row) and gastric adenocarcinoma^34^ (GAC, bottom row). Violin plots display gene expression levels in fibroblasts from adjacent-normal (blue) and tumor (red) tissues.

## References

1. Tlsty, T. D. & Gascard, P. Stromal directives can control cancer. Science 365, 122–123 (2019).

2. Lau, S. C. M., Pan, Y., Velcheti, V. & Wong, K. K. Squamous cell lung cancer: Current landscape and future therapeutic options. Cancer Cell 40, 1279–1293 (2022).

3. Chen, Q., Zheng, X., Cheng, W. & Li, J. Landscape of targeted therapies for lung squamous cell carcinoma. Front. Oncol. 14, 1467898 (2024).

4. Niu, Z., Jin, R., Zhang, Y. & Li, H. Signaling pathways and targeted therapies in lung squamous cell carcinoma: mechanisms and clinical trials. Signal Transduct. Target. Ther. 7, 353 (2022).

5. Wang, Y. et al. Immunotherapy for advanced-stage squamous cell lung cancer: the state of the art and outstanding questions. Nat. Rev. Clin. Oncol. 22, 200–214 (2025).

6. Walser, T., et al. Smoking and Lung Cancer. Proc. Am. Thorac. Soc. 5, 811–815 (2008).

7. Sharma, V., Letson, J. & Furuta, S. Fibrous stroma: Driver and passenger in cancer development. Sci. Signal. 15, eabg3449 (2022).

8. Bulle, A. & Lim, K.-H. Beyond just a tight fortress: contribution of stroma to epithelial-mesenchymal transition in pancreatic cancer. Signal Transduct. Target. Ther. 5, 249 (2020).

9. Chandler, C., Liu, T., Buckanovich, R. & Coffman, L. G. The double edge sword of fibrosis in cancer. Transl. Res. 209, 55–67 (2019).

10. Yamauchi, M., Barker, T. H., Gibbons, D. L. & Kurie, J. M. The fibrotic tumor stroma. J. Clin. Investig. 128, 16–25 (2018).

11. Pan, D. et al. A Fibroblast State Choreographs an Epithelial YAP-dependent Regenerative Program Essential to (Pre)malignancy via ECM-mediated Mechanotransduction. bioRxiv 2025.07.11.661192 (2025) doi:10.1101/2025.07.11.661192.

12. Adler, M. et al. Principles of *Cell* Circuits for Tissue Repair and Fibrosis. Iscience 23, 100841 (2020).

13. Alon, U. Systems Medicine, Physiological Circuits and the Dynamics of Disease. (2023) doi:10.1201/9781003356929.

14. Miyara, S. et al. Cold and hot fibrosis define clinically distinct cardiac pathologies. Cell Syst. 101198 (2025) doi:10.1016/j.cels.2025.101198.

15. Setten, E. et al. Understanding fibrosis pathogenesis via modeling macrophage-fibroblast interplay in immune-metabolic context. Nat Commun 13, 6499 (2022).

16. Zhou, X. et al. Circuit Design Features of a Stable Two-Cell System. Cell 172, 744–757.e17 (2018).

17. Wang, S. et al. An autocrine signaling circuit in hepatic stellate cells underlies advanced fibrosis in nonalcoholic steatohepatitis. Sci Transl Med 15, eadd3949 (2023).

18. Hart, Y. et al. Inferring biological tasks using Pareto analysis of high-dimensional data. Nat Methods 12, 233–235 (2015).

19. Mayer, S. et al. The tumor microenvironment shows a hierarchy of cell-cell interactions dominated by fibroblasts. Nat. Commun. 14, 5810 (2023).

20. Lizotte, P. H., et al. Multiparametric profiling of non–small-cell lung cancers reveals distinct immunophenotypes. JCI Insight 1, e89014 (2016).

21. Zhang, H. et al. Spatial positioning of immune hotspots reflects the interplay between B and T cells in lung squamous cell carcinoma. Cancer Res. 83, 1410–1425 (2023).

22. Enfield, K. S. S. et al. Spatial Architecture of Myeloid and T Cells Orchestrates Immune Evasion and Clinical Outcome in Lung Cancer. Cancer Discov. 14, 1018–1047 (2024).

23. Lawrence, M. S. et al. Comprehensive genomic characterization of head and neck squamous cell carcinomas. Nature 517, 576–582 (2015).

24. Watanabe, H. et al. SOX2 and p63 colocalize at genetic loci in squamous cell carcinomas. J. Clin. Investig. 124, 1636–1645 (2014).

25. Sauler, M. et al. Characterization of the COPD alveolar niche using single-cell RNA sequencing. Nat. Commun. 13, 494 (2022).

26. Patir, A. et al. The transcriptional signature associated with human motile cilia. Sci. Rep. 10, 10814 (2020).

27. Lavie, D., Ben-Shmuel, A., Erez, N. & Scherz-Shouval, R. Cancer-associated fibroblasts in the single-cell era. *Nat*. Cancer 3, 793–807 (2022).

28. Torregrossa, M. et al. Effects of embryonic origin, tissue cues and pathological signals on fibroblast diversity in humans. Nat. Cell Biol. 1–16 (2025) doi:10.1038/s41556-025-01638-5.

29. Davidson, G. et al. Mesenchymal-like Tumor Cells and Myofibroblastic Cancer-Associated Fibroblasts Are Associated with Progression and Immunotherapy Response of Clear Cell Renal Cell Carcinoma. Cancer Res. 83, 2952–2969 (2023).

30. Barkley, D. et al. Cancer cell states recur across tumor types and form specific interactions with the tumor microenvironment. Nat. Genet. 54, 1192–1201 (2022).

31. Peters, Y., et al. Barrett oesophagus. Nat. Rev. Dis. Prim. 5, 35 (2019).

32. Thrift, A. P., Wenker, T. N. & El-Serag, H. B. Global burden of gastric cancer: epidemiological trends, risk factors, screening and prevention. Nat. Rev. Clin. Oncol. 20, 338–349 (2023).

33. Strasser, M. K. et al. Concerted changes in Epithelium and Stroma: a multi-scale, multi-omics analysis of progression from Barrett’s Esophagus to adenocarcinoma. Dev. Cell (2025) doi:10.1016/j.devcel.2025.06.034.

34. Kumar, V. et al. Single-Cell Atlas of Lineage States, Tumor Microenvironment, and Subtype-Specific Expression Programs in Gastric CancerSingle-Cell Atlas of Gastric Cancer Subtypes. Cancer Discov. 12, 670–691 (2022).

35. Browaeys, R., Saelens, W. & Saeys, Y. NicheNet: modeling intercellular communication by linking ligands to target genes. Nat Methods 17, 159–162 (2020).

36. Aegerter, H., Lambrecht, B. N. & Jakubzick, C. V. Biology of lung macrophages in health and disease. Immunity 55, 1564–1580 (2022).

37. Sahai, E. et al. A framework for advancing our understanding of cancer-associated fibroblasts. Nat. Rev. Cancer 20, 174–186 (2020).

38. Friedman, G. et al. Cancer-associated fibroblast compositions change with breast cancer progression linking the ratio of S100A4+ and PDPN+ CAFs to clinical outcome. Nat Cancer 1, 692–708 (2020).

39. Quail, D. F. & Joyce, J. A. Microenvironmental regulation of tumor progression and metastasis. Nat. Med. 19, 1423–1437 (2013).

40. Flier, J. S., Underhill, L. H. & Dvorak, H. F. Tumors: Wounds That Do Not Heal. N. Engl. J. Med. 315, 1650–1659 (1986).

41. Kikuchi, K., Kadono, T., Furue, M. & Tamaki, K. Tissue Inhibitor of Metalloproteinase 1 (TIMP-1) May Be an Autocrine Growth Factor in Scleroderma Fibroblasts. J Invest Dermatol 108, 281–284 (1997).

42. Fowell, A. J. et al. Silencing tissue inhibitors of metalloproteinases (TIMPs) with short interfering RNA reveals a role for TIMP-1 in hepatic stellate cell proliferation. Biochem Bioph Res Co 407, 277–282 (2011).

43. Bertaux, B., Hornebeck, W., Eisen, A. Z. & Dubertret, L. Growth Stimulation of Human Keratinocytes by Tissue Inhibitor of Metalloproteinases. J Invest Dermatol 97, 679–685 (1991).

44. Hayakawa, T., Yamashita, K., Tanzawa, K., Uchijima, E. & Iwata, K. Growth-promoting activity of tissue inhibitor of metalloproteinases-1 (TIMP-1) for a wide range of cells A possible new growth factor in serum. FEBS Lett. 298, 29–32 (1992).

45. Schoeps, B., Frädrich, J. & Krüger, A. Cut loose TIMP-1: an emerging cytokine in inflammation. Trends Cell Biol. 33, 413–426 (2023).

46. Aoki, F., Kurabayashi, M., Hasegawa, Y. & Kojima, I. Attenuation of Bleomycin-induced Pulmonary Fibrosis by Follistatin. Am. J. Respir. Crit. Care Med. 172, 713–720 (2005).

47. Grunberg, N. et al. Cancer-Associated Fibroblasts Promote Aggressive Gastric Cancer Phenotypes via Heat Shock Factor 1–Mediated Secretion of Extracellular Vesicles. Cancer Res. 81, 1639–1653 (2021).

48. Fordyce, C. A. et al. Cell-extrinsic consequences of epithelial stress: activation of protumorigenic tissue phenotypes. Breast Cancer Res. 14, R155 (2012).

49. Frangogiannis, N. G. Transforming growth factor–β in tissue fibrosis. J. Exp. Med. 217, e20190103 (2020).

50. Flavell, R. A., Sanjabi, S., Wrzesinski, S. H. & Licona-Limón, P. The polarization of immune cells in the tumour environment by TGFβ. Nat. Rev. Immunol. 10, 554–567 (2010).

51. Batlle, E. & Massagué, J. Transforming Growth Factor-β Signaling in Immunity and Cancer. Immunity 50, 924–940 (2019).

52. Mariathasan, S. et al. TGFβ attenuates tumour response to PD-L1 blockade by contributing to exclusion of T cells. Nature 554, 544–548 (2018).

53. Fan, J. et al. Glia maturation factor-β: a potential therapeutic target in neurodegeneration and neuroinflammation. Neuropsychiatr. Dis. Treat. 14, 495–504 (2018).

54. Sun, W. et al. Glia Maturation Factor Beta as a Novel Biomarker and Therapeutic Target for Hepatocellular Carcinoma. Front. Oncol. 11, 744331 (2021).

55. Sun, R. et al. Amphiregulin couples IL1RL1+ regulatory T cells and cancer-associated fibroblasts to impede antitumor immunity. Sci. Adv. 9, eadd7399 (2023).

56. Cook, S. A. & Schafer, S. Hiding in Plain Sight: Interleukin-11 Emerges as a Master Regulator of Fibrosis, Tissue Integrity, and Stromal Inflammation. Annu. Rev. Med. 71, 263–276 (2020).

57. Feig, C. et al. Targeting CXCL12 from FAP-expressing carcinoma-associated fibroblasts synergizes with anti–PD-L1 immunotherapy in pancreatic cancer. Proc. Natl. Acad. Sci. 110, 20212–20217 (2013).

58. Chitty, J. L. et al. A first-in-class pan-lysyl oxidase inhibitor impairs stromal remodeling and enhances gemcitabine response and survival in pancreatic cancer. Nat. Cancer 1–19 (2023) doi:10.1038/s43018-023-00614-y.

59. Nielsen, S. R. et al. Macrophage-secreted granulin supports pancreatic cancer metastasis by inducing liver fibrosis. Nat. Cell Biol. 18, 549–560 (2016).

60. Watson, S. S. et al. Fibrotic response to anti-CSF-1R therapy potentiates glioblastoma recurrence. Cancer Cell 42, 1507–1527.e11 (2024).

61. Özdemir, B. C. et al. Depletion of Carcinoma-Associated Fibroblasts and Fibrosis Induces Immunosuppression and Accelerates Pancreas Cancer with Reduced Survival. Cancer Cell 25, 719–734 (2014).

62. Aronesty, E. Comparison of Sequencing Utility Programs. Open Bioinform. J. 7, 1–8 (2013).

63. Martin, M. Cutadapt removes adapter sequences from high-throughput sequencing reads. Embnet J 17, 10–12 (2011).

64. Bray, N. L., Pimentel, H., Melsted, P. & Pachter, L. Near-optimal probabilistic RNA-seq quantification. Nat. Biotechnol. 34, 525–527 (2016).

65. Melsted, P. et al. Modular, efficient and constant-memory single-cell RNA-seq preprocessing. Nat. Biotechnol. 39, 813–818 (2021).

66. Wolf, F. A., Angerer, P. & Theis, F. J. SCANPY: large-scale single-cell gene expression data analysis. Genome Biol. 19, 15 (2018).

67. Wolock, S. L., Lopez, R. & Klein, A. M. Scrublet: Computational Identification of Cell Doublets in Single-Cell Transcriptomic Data. Cell Syst. 8, 281–291.e9 (2019).

68. Zheng, G. X. Y. et al. Massively parallel digital transcriptional profiling of single cells. Nat. Commun. 8, 14049 (2017).

69. Korsunsky, I. et al. Fast, sensitive and accurate integration of single-cell data with Harmony. Nat. Methods 16, 1289–1296 (2019).

70. Wolf, F. A. et al. PAGA: graph abstraction reconciles clustering with trajectory inference through a topology preserving map of single cells. Genome Biol. 20, 59 (2019).

71. Traag, V. A., Waltman, L. & Eck, N. J. van. From Louvain to Leiden: guaranteeing well-connected communities. Sci. Rep. 9, 5233 (2019).

72. Satija, R., Farrell, J. A., Gennert, D., Schier, A. F. & Regev, A. Spatial reconstruction of single-cell gene expression data. Nat. Biotechnol. 33, 495–502 (2015).

73. Butler, A., Hoffman, P., Smibert, P., Papalexi, E. & Satija, R. Integrating single-cell transcriptomic data across different conditions, technologies, and species. Nat. Biotechnol. 36, 411–420 (2018).

74. Breda, J., Zavolan, M. & Nimwegen, E. van. Bayesian inference of gene expression states from single-cell RNA-seq data. Nat Biotechnol 39, 1008–1016 (2021).

75. Bonnardel, J., et al. Stellate Cells, Hepatocytes, and Endothelial Cells Imprint the Kupffer Cell Identity on Monocytes Colonizing the Liver Macrophage Niche. Immunity 51, 638–654.e9 (2019).

76. Guilliams, M. et al. Spatial proteogenomics reveals distinct and evolutionarily conserved hepatic macrophage niches. Cell 185, 379–396.e38 (2022).

